# The genome of low-chill Chinese plum ‘Sanyueli’ (*Prunus salicina* Lindl.) provides insights into the regulation of the chilling requirement of flower buds

**DOI:** 10.1101/2020.07.31.193243

**Authors:** Zhi-Zhen Fang, Kui Lin-Wang, He Dai, Dan-Rong Zhou, Cui-Cui Jiang, Richard V. Espley, Yan-Juan Lin, Shao-Lin Pan, Xin-Fu Ye

**Affiliations:** Fruit Research Institute, Fujian Academy of Agricultural Sciences, Fuzhou, Fujian 350013, China; Fujian Engineering and Technology Research Center for Deciduous Fruit Trees, Fujian Academy of Agricultural Sciences, Fuzhou, Fujian 350013, China; The New Zealand Institute for Plant and Food Research Limited, Mt Albert Research Centre, Private Bag, Auckland 92169, New Zealand; Biomarker Technologies Corporation, Beijing 101300, China

**Author notes:** Corresponding author, Fruit Research Institute, Fujian Academy of Agricultural Sciences, Hualin Road 247, Fuzhou 350003, China.

## Abstract

Chinese plum (*Prunus salicina* Lindl.) is a stone fruit that belongs to the *Prunus* genus and plays an important role in the global production of plum. In this study, we report the genome sequence of the Chinese plum ‘Sanyueli’, which is known to have a low-chill requirement for flower bud break. The assembled genome size was 308.06 Mb, with a contig N50 of 815.7 kb. A total of 30,159 protein-coding genes were predicted from the genome and 56.4% (173.39 Mb) of the genome was annotated as repetitive sequence. Bud dormancy is influenced by chilling requirement in plum and partly controlled by *DORMANCY ASSOCIATED MADS-box* (*DAM*) genes. Six tandemly arrayed *PsDAM* genes were identified in the assembled genome. Sequence analysis of *PsDAM6* in ‘Sanyueli’revealed the presence of large insertions in the intron and exon regions. Transcriptome analysis indicated that the expression of *PsDAM6* in the dormant flower buds of ‘Sanyueli’ was significantly lower than that in the dormant flower buds of the high chill requiring ‘Furongli’ plum. In addition, the expression of *PsDAM6* was repressed by chilling treatment. The genome sequence of ‘Sanyueli’ plum provides a valuable resource for elucidating the molecular mechanisms responsible for the regulation of chilling requirements, and is also useful for the identification of the genes involved in the control of other important agronomic traits and molecular breeding in plum.

## Introduction

Plums are temperate fruit trees, which belong to *Prunus* genus in the *Rosaceae* family, and have been grown throughout the world for centuries. Plums provide the second largest stone fruit production after peaches and nectarines. Chinese plum (also known as Japanese plum, *Prunus salicina* Lindl.) and European plum (*Prunus domestica* L.) are the most commercially significant species^1^. Chinese plum is native to China^1^. The discovery of fossil plum stone dated to the Neolithic or the Warring States period implicated that plum fruits have been used as food for over 5,000-6,000 years in China ^2^. The discoveries of plum stones in tombs of ancient Chinese people and an in-depth description of plum in the Book of Odes^2,3^, a collection of poems dating from 1,100 BC to 600 BC, suggested that plum has been cultivated and popular in China for over 3,000 years. Chinese plum was transported to Japan over 2,000 years ago and was introduced into the USA in the 1870s^1,4^. It was estimated that Chinese plum and its hybrids account for over 70% of the world’s global plum production^1^. China is the largest plum producer in the world and almost all of the production is Chinese plum^1^. According to the Food and Agriculture Organization of the United Nations (FAO), plum fruit production in China was over 680 million tons in 2017.

Similar to other temperate/deciduous fruit trees, plum trees require a certain amount of cool temperature during the dormancy period to fulfill their chilling requirements and allow for the release of dormancy. Most Chinese plum varieties require 500-800 hours of chilling and European plums need even more (often >1,000 hours)^5^. Lack of winter chill results in delayed and abnormal flowering and negatively affects the yield and quality of fruits^6,7^. Plums are mainly grown in temperate zones^8^. In China, the production of plum is mainly concentrated in subtropical regions, including Guangdong, Guangxi and Fujian^9^. In the context of global warming, it is particularly pressing to develop low-chill plum cultivars^8,10^. A low chilling requirement has long been one of the principal objectives in plum breeding and breeding programs to develop low-chill cultivars are evident in many countries, especially in the southern hemisphere^1,4,11–13^. Understanding the molecular mechanisms responsible for the regulation of bud dormancy and chilling requirements in plums is essential for the breeding of low-chill cultivars and ensuring consistent plum production in a changing environment.

Bud dormancy has attracted a great amount of attention in recent decades due to the agronomic disorders caused by warm winters^14–16^. Considerable efforts have been made to uncover the genetic basis of bud dormancy in Rosaceae species, including apple^17–24^, peach^25–35^, mume^36–41^, pear^42–50^, sweet cherry^51–54^, almond^55,56^ and apricot^57–61^. These studies as well as the publications on other species have emphasized the role of *DORMANCY ASSOCIATED MADS-box* (*DAM*) genes in regulation of bud dormancy and shed light on the transcriptional regulation of these genes^14^. However, to date the mechanisms involved in modulation of bud dormancy and chilling requirement in plum is still unknown.

To date the genome sequences of several *Prunus* species of the *Rosaceae* family, including mume^62^, peach^63^, sweet cherry^64^, flowering cherry^65^, almond^66,67^, apricot^68^, have been published. Recently, the genome sequence of European plum has become available in the Genome Database for Rosaceae^69^. Genomes of commercial crops are valuable resources for the identification of genetic loci regulating important agronomic traits and molecular breeding in the future. The availability of these genomes has enabled the identification of genes for certain traits^70^. Currently, genome information for Chinese plum and its hybrids is not available. In this study, we sequenced and assembled the genome of Chinese plum ‘Sanyueli’, a low-chilling requirement cultivar. He, et al.^71^ reported that ‘Sanyueli’ has the lowest chilling requirement among the varieties cultivated in Guangzhou, Guangdong Province, China. Later, they demonstrated that 28 h of chilling temperatures (≤7.2◻) was sufficient to fulfill the chilling requirement of ‘Sanyueli’ flower buds^72^. We also conducted transcriptomic analyses to identify candidate genes underlying chilling requirement in plum. The results extend our understanding of the molecular control of dormancy and chilling requirement in plum. This genome sequence will help facilitate genetic research of many agronomic traits support the development of novel plum cultivars.

## Results

### Genome assembly

‘Sanyueli’ is an early-maturing and high-yielding plum variety with red skin and yellow flesh (Fig. 1A-C). ‘Sanyueli’ trees are able to bloom and fruit without exposure to a large number of chilling hours (Fig. 1D-E). Our results showed that the chilling requirement of ‘Sanyueli’ flower buds was fulfilled after 50 h at chilling temperature (Supplementary Fig. 2A). However, nearly 800 h was required for breaking the flower bud dormancy of ‘Furongli’ (Supplementary Fig. 2B).

**Fig. 1.**
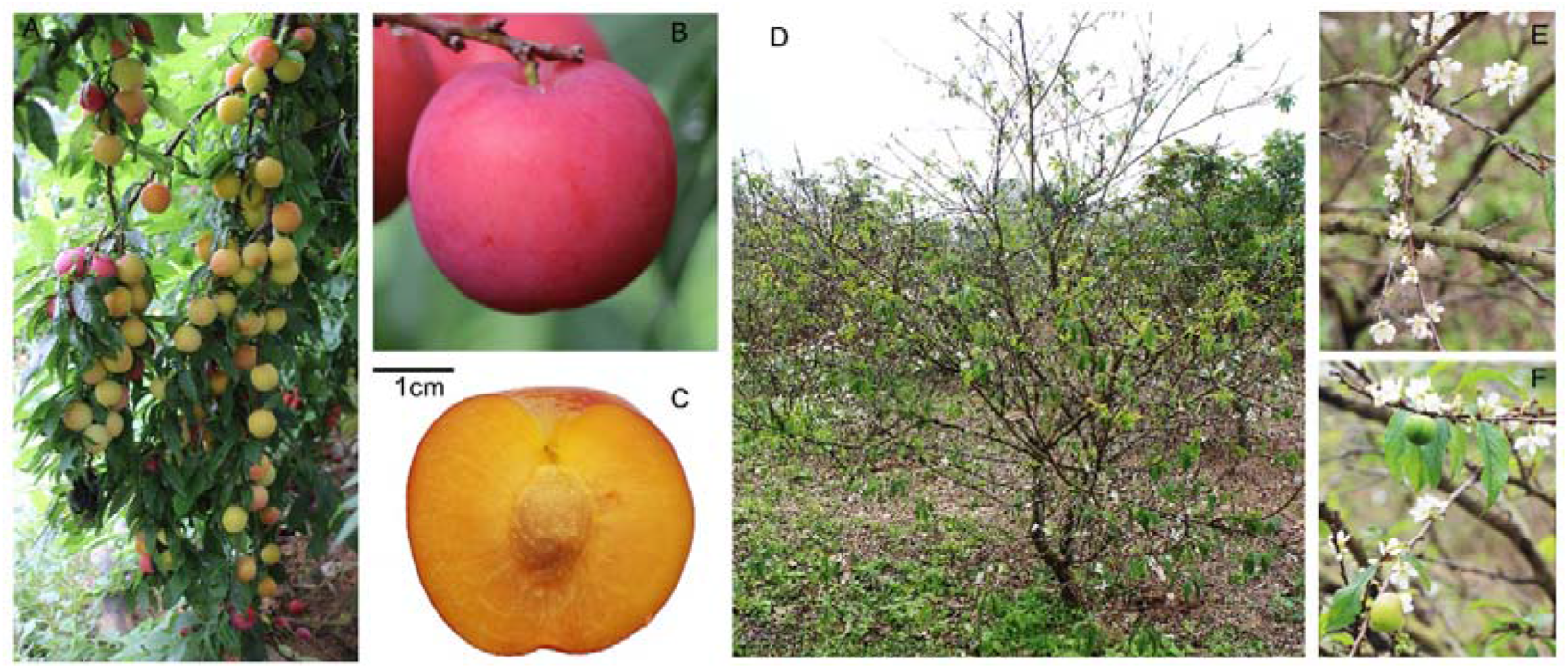
Images of ‘Sanyueli’ plum used in this study. A, B and C indicate fruit of ‘Sanyueli’. D shows a ‘Sanyueli’ plum tree with flowers (E) and fruits (F) in Zhangpu county (Zhangzhou, Fujian province, China) on 5 February, 2016. Daily maximum and minimum temperature from November 2015 to February 2016 was indicated in Supplementary Fig. 1.

We sequenced and assembled the genome of ‘Sanyueli’ using a combination of short-read sequencing from Illumina HiSeq 4000 and SMRT from Pacific Biosciences (PacBio, Menlo Park, CA). First, we generated 28.08 Gb Illumina paired-end (PE) reads with insert sizes of 270 bp (Supplementary Table 1) and used them to estimate the genome size and heterozygosity ratio of ‘Sanyueli’ genome. Based on a 19-mer analysis, we evaluated the genome size to be 308.06 Mb, with a heterozygosity of 0.33%, and the estimated repeat sequence content being 52.6% (Supplementary Fig.3). Then 14.89 Gb of PacBio reads (Supplementary Table 2) were produced and assembled into 1,752 contigs (longest being 7,898,106 bp) with an N50 of 815.7 kb. The size of the final assembled genome was 307.29 Mb, which is very close to the estimated size. The GC content of the assembled plum genome was 37.8% (Table 1).

**Table 1.** Statistics of ‘Sanyueli’ plum genome assembly and annotation.

|  |  |
| --- | --- |
| Genome size (Mb) | 307.29 |
| Number of contigs ($\geq 1$ kb) | 1,752 |
| N50 contig length (bp) | 815,770 |
| N90 contig length (bp) | 50,776 |
| Largest contig (bp) | 7,898,106 |
| GC content (%) | 37.78 |
| Number of gene models | 30,159 |
| Gene length (bp) | 108,356,597 |
| Mean gene length (bp) | 3592.8 |
| Total coding region length (bp) | 39,130,040 |
| Mean coding region length (bp) | 1297.4 |
| Total intron length (bp) | 58,765,475 |
| Mean intron length (bp) | 437.5 |

The quality of the assembled genome was evaluated using three strategies. First, BWA alignment results indicated that 97.9% of the Illumina paired-end reads were successfully aligned to the genome (Supplementary Table 1). Second, according to BUSCO (version 2)^73^, our assembly contained 97.0% (1397 of 1440) of the core eukaryotic genes, including 1292 single-copy orthologs and 197 duplicated orthologs (Supplementary Table 3). Third, CEGMA^74^ results indicated that 451 out of 458 highly conserved core genes were detected in our assembly (Supplementary Table 4), suggesting the high completeness of our genome assembly. These results indicated that our plum genome sequence was almost complete.

### Genome annotation

#### Annotation of repeat sequences

56.4% (173.39 Mb) of the assembled genome was predicted to be repetitive, which is higher than the repeat content observed in mume (44.9%), sweet cherry (43.8%), apricot (38.3%), peach (37.1%) and almond (34.6%). The repetitive sequences including retrotransposons (Class I elements, 48.34%), DNA transposons (Class II elements, 12.0%), potential host genes (1.6%), simple sequence repeats (0.02%) and unclassified elements (3.8%) (Supplementary Table 5). The proportion of Gypsy retrotransposon (24.4%) appears to have expanded considerably in the ‘Sanyueli’ genome compared with that of the peach (10.0%) and mume (8.6%) and is comparable with that of apple (25.2%). The proportion of full-length long terminal repeats (LTR)/Copia repeats (9.3%) in the ‘Sanyueli’ genome was similar to that in mume (10.0%) and higher than that in peach (8.6%) and apple (5.0%). The PLE/LARD retrotransposon derivative repeat elements represented 9.6% of the genome, similar to the proportion of the genome represented by LTR/ Copia elements (Supplementary Table 5).

#### Gene prediction and functional annotation

A combination of *ab initio*, homology-based and RNA-seq based prediction methods was used to predict gene models from the plum genome sequence. A total of 30,159 protein-coding genes were predicted (representing 35.3% of the genome assembly), with an average gene length of 3593 bp, and an average coding region sequence size of 1297 bp (Supplementary Table 6). The average gene density of plum was 98 genes per Mb, which is lower than in apricot (137 genes per Mb), mume (132 genes per Mb), peach (122 genes per Mb) and almond (112 genes per Mb), but is higher than in sweet cherry (87 genes per Mb). 21,939 genes (72.7%) were supported by RNA-seq data and 29,190 genes (96.8%) were supported by homology to known proteins. A total of 19,486 genes (64.6%) were supported by all three methods (Supplementary Fig. 4), and these genes were annotated with high confidence. A total of 29,817 (98.87%) protein-coding genes were annotated based on GO, Kyoto Encyclopedia of Genes and Genomes (KEGG), EuKaryotic Orthologous Groups (KOG), Pfam, Swissprot, TrEMBL, Nr and Nt databases (Supplementary Table 7). In addition, 85 microRNAs (miRNAs), 602 ribosomal RNAs, 525 transfer RNAs, 2210 small nucleolar RNAs (snoRNAs), 101 small nuclear RNAs (snRNAs) and 1419 pseudogenes were identified in the ‘Sanyueli’ plum genome (Supplementary Table 8).

#### Comparative genomic and genome evolutionary analysis

We performed gene family cluster analysis on the genomes of *P. salicina*, *P. armeniaca*, *P. persica*, *P. avium*, *P. mume*, *P. dulcis*, *P. bretschneideri*, *M.* × *domestica*, *F. vesca*, *R. chinensis*, *V. vinifera*, *P. trichocarpa*, *O. sativa*, *A. thaliana* and *A. trichopoda.* The number of single-copy genes in *P. salicina* was lower than in other *Prunus* species, but higher than in *P. bretschneideri* and *M. × domestica*, while the number of multi-copy genes in *P. salicina* was higher than in other *Prunus* species and lower than in *P. bretschneideri* and *M.* × *domestica* (Fig. 2A). A total of 28006 genes from the plum genome were grouped into 17622 gene clusters and 219 gene families containing 553 genes (Supplementary Table 9 and 10) were identified as plum specific. We compared the gene numbers among the six *Prunus* species. 11053 gene families were shared by all *Prunus* species, and 288 gene families were specific to *P. salicina*, which is higher than the number found in genomes *P. persica* and *P. mume*, but lower than those of *P.avium, P.armeniaca* and *P.dulcis* (Fig. 2B).

**Fig. 2.**
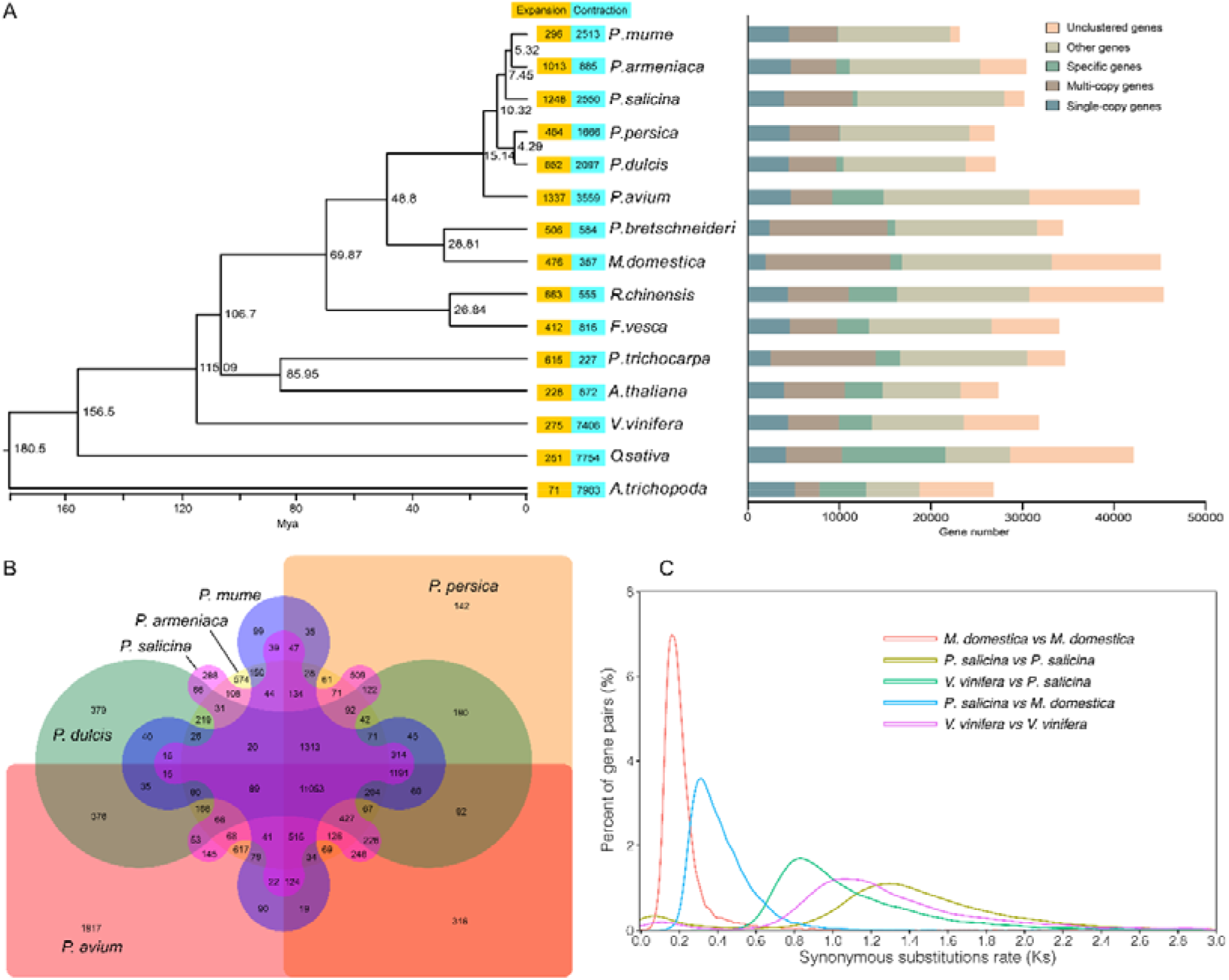
Comparative genomic analysis of plum and other species. A, Phylogenetic analysis and gene family cluster analysis of plum and 14 other species. The phylogenetic tree (left panel) was constructed using 436 single-copy genes from plum and 14 other species, including *Prunus armeniaca*, *P. persica*, *P. avium*, *P. mume*, *P. dulcis*, *Pyrus bretschneideri, Malus* × *domestica*, *Fragaria vesca*, *Rosa chinensis*, *Vitis vinifera*, *Populus trichocarpa*, *Oryza sativa*, *Arabidopsis thaliana* and *Amborella trichopoda.* The numbers indicate the divergence time. Gene family expansions and contractions are indicated by numbers in the yellow and blue boxes. The distribution of paralogous genes in the analyzed plants is indicated in the right panel. B, Venn diagram comparison of gene families in six *Prunus* species. C, Distribution of synonymous nucleotide substitutions (Ks) among *P. salicina*, *M.* × *domestica* and *V. vinifera*. Duplication components were identified by MCScan without tandem duplicated pairs. Colored curves superimposed on the Ks distribution represent different components identified. Ks distributions of paralogous gene pairs were identified from syntenic blocks between species, and orthologous gene pairs were identified from duplication components within one genome.

A phylogenetic tree of plum, five *Prunus* species and nine other sequenced species was constructed using single-copy genes. The phylogenetic tree showed that *P. salicina* is relatively closely related to *P. mume* and *P. armeniaca* (Fig. 2A). It also indicated that *P. salicina* diverged from *P. mume* and *P. armeniaca* approximately 7.45 million years ago (Mya) and from *P. dulcis* and *P. persica* 10.32 Mya, after the divergence of *P. avium* at 15.14 Mya (Fig. 2A). Genome collinearity analysis demonstrated that the*P. salicina* genome showed high collinear relationships with the genome of *P. armeniaca* and *P. mume*, which are closely related to *P. salicina*. (Supplementary Fig. 5).

The synonymous nucleotide substitution (Ks) analysis in plum, *M.* × *domestica* and *V. vinifera* indicated that there has been an ancient WGD (ancient γ whole-genome duplication) in the plum genome before the divergence with *M.* × *domestica* and *V. vinifera* (Fig. 2C). In addition, 1248 and 2550 gene families were found to have expanded and contracted in the plum genome, which is more than the number of expanded and contracted gene families in the genomes of *P. mume* and *P.armeniaca*. KEGG enrichment analysis of the expanded genes showed that plant-pathogen interaction, starch and sucrose metabolism and phenylpropanoid biosynthesis were the most enriched pathways (Supplementary Table 11). Among the 74 genes assigned to the starch and sucrose metabolism pathway, 17 were annotated as polygalacturonase and two of them (PsSY0006672 and PsSY0008876) were significantly upregulated in the flesh during fruit ripening. GO analysis of the expanded orthogroups revealed that the oxidation-reduction process (GO:0055114), regulation of transcription, DNA-templated (GO:0006355), metabolic process (GO:0008152) and protein phosphorylation (GO:0006468) were significantly enriched (Supplementary Table 12).

#### *DORMANCY ASSOCIATED MADS-box* (*DAM*) genes in plum genome

*DAM* genes have been reported to be the major regulatory factors involved in the control of dormancy induction and release in flower buds of *Prunus* species^61,75,76^. Blast analyses against the genome of ‘Sanyueli’ plum enabled us to identify six *PsDAM* genes, including *PsDAM1* (PsSY0028611), *PsDAM2* (PsSY0012373), *PsDAM3* (PsSY0007206), *PsDAM4* (PsSY0000114), *PsDAM5* (PsSY0007447) and *PsDAM6* (PsSY0011722), which are tandemly arrayed in the genome of ‘Sanyueli’ plum (Fig. 3A). Genomic structure analysis revealed that *PsDAM1*, *PsDAM2*, *PsDAM3*, *PsDAM4* and *PsDAM5* have genomic structures consisting of eight exons and seven introns similar to their homologs from other *Prunus* species (Fig. 3B). However, *PsDAM6* has only six exons, which was different from *DAM6* genes in other *Prunus* species. In addition, *PsDAM6* is much longer because of the large size of the fourth intron, when compared with other *Prunus* species (Fig. 3B). Sequence analysis of *PsDAM6* gene from ‘Sanyueli’ and ‘Furongli’ plums indicated there were several insertions, including two large insertions (1327bp and 6493bp) (Fig. 3C and Supplementary Fig. 6). The 6493bp insertion was detected in exon5 of *PsDAM6* from ‘Sanyueli’, which causes the loss of function of exon5 (Fig. 3C and Supplementary Fig. 6). Furthermore, a fragment in the seventh intron of *PsDAM6* from ‘Furongli’ was reverse-inserted into the first intron of the *PsDAM6* genes from ‘Sanyueli’ (Fig. 3C and Supplementary Fig. 6). Phylogenetic analysis indicated that PsDAMs group into a clade with DAMs from other *Prunus* species and the *DAM1-DAM6* sequences formed a subgroup with the respective sequences from other *Prunus* species (Fig. 4). Multiple sequence alignment of plum PsDAMs with DAMs from Prunus species indicated that all *Prunus* DAM proteins contain a conserved MADS domain, an I domain and a K domain(Fig. 5). A conserved EAR motif, which acts as a repression domain, was also detected in the C-terminal of these DAM proteins (Fig. 5). The deletion of exon5 led to the lack of 14 amino

**Fig. 3.**
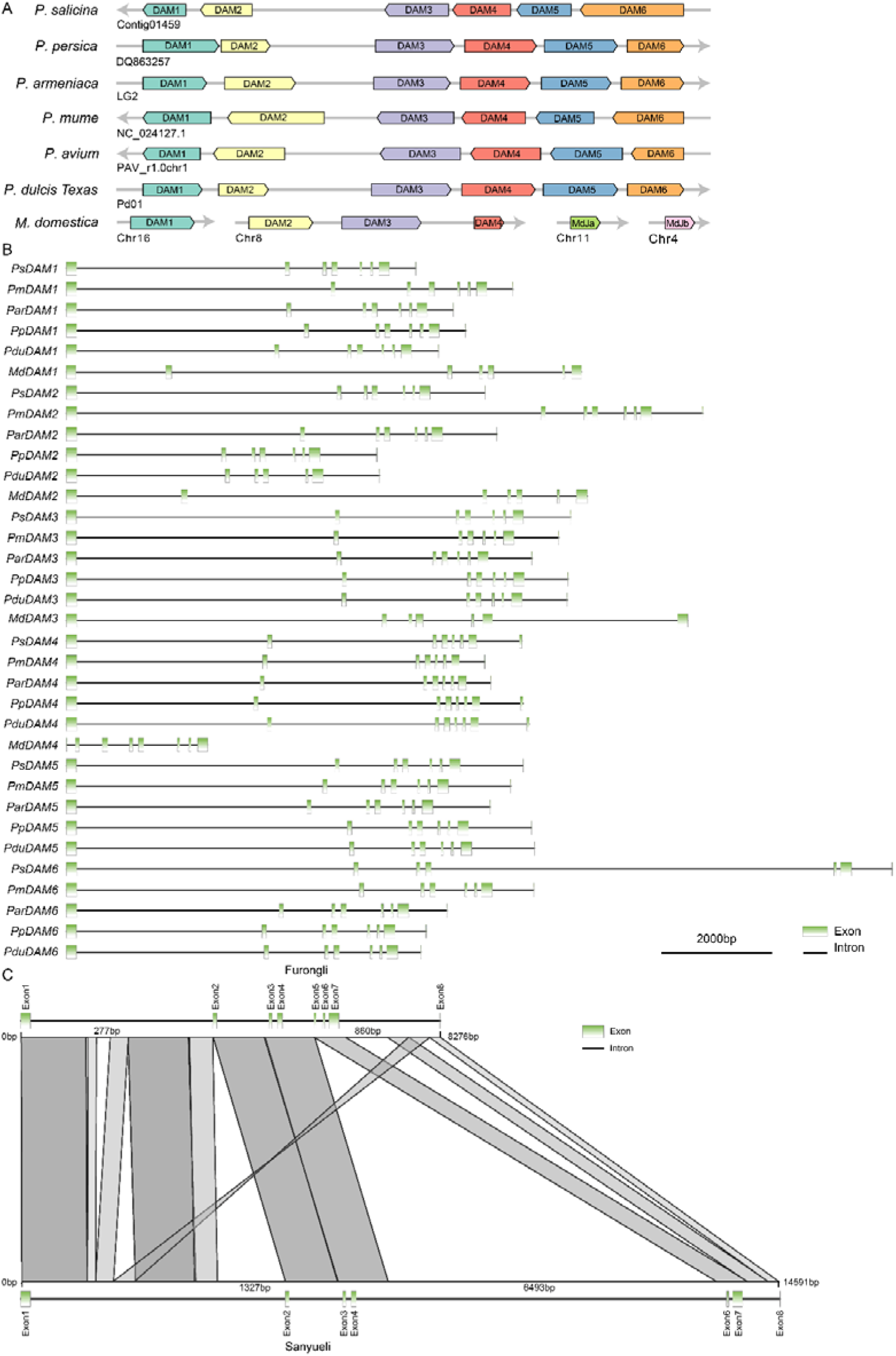
*PsDAM* genes in plum and other Rosaceae species. A, Overview of the *DAM* locus in the genome of ‘Sanyueli’ plum and other Rosaceae species. *DAM* genes showing synteny are indicated in the same color. B, Schematic overview of introns and exons in *DAM* genes from plum and other Rosaceae species. C, Boxes and lines represent exons and introns, respectively. Structural alignments of the *PsDAM6* gene from ‘Sanyueli’ and ‘Furongli’ plum. acid residues, which belong to K3 helix in the K domain (Fig. 5). These results suggested that DAMs are conserved in *Prunus* species and plum PsDAMs may have similar function with their homologs from *Prunus* species.

**Fig. 4.**
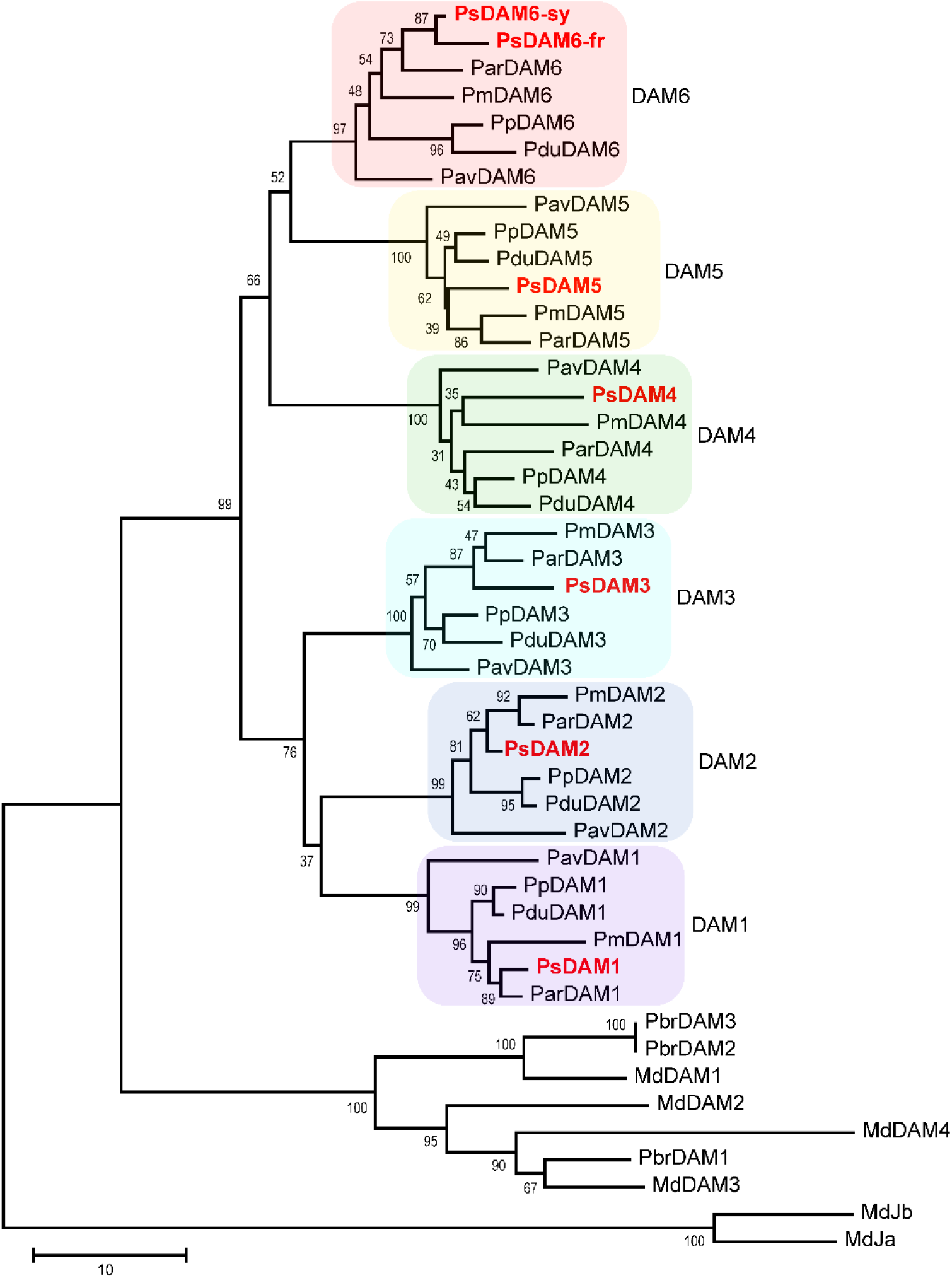
Phylogenetic analysis of predicted amino acid sequences of *DAM* genes from plum and other Rosaceae species. DAMs in plum are colored by red. PsDAM6-sy and PsDAM6-fr represent PsDAM6 from ‘Sanyueli’ and ‘Furongli’ plum, respectively.

**Fig. 5.**
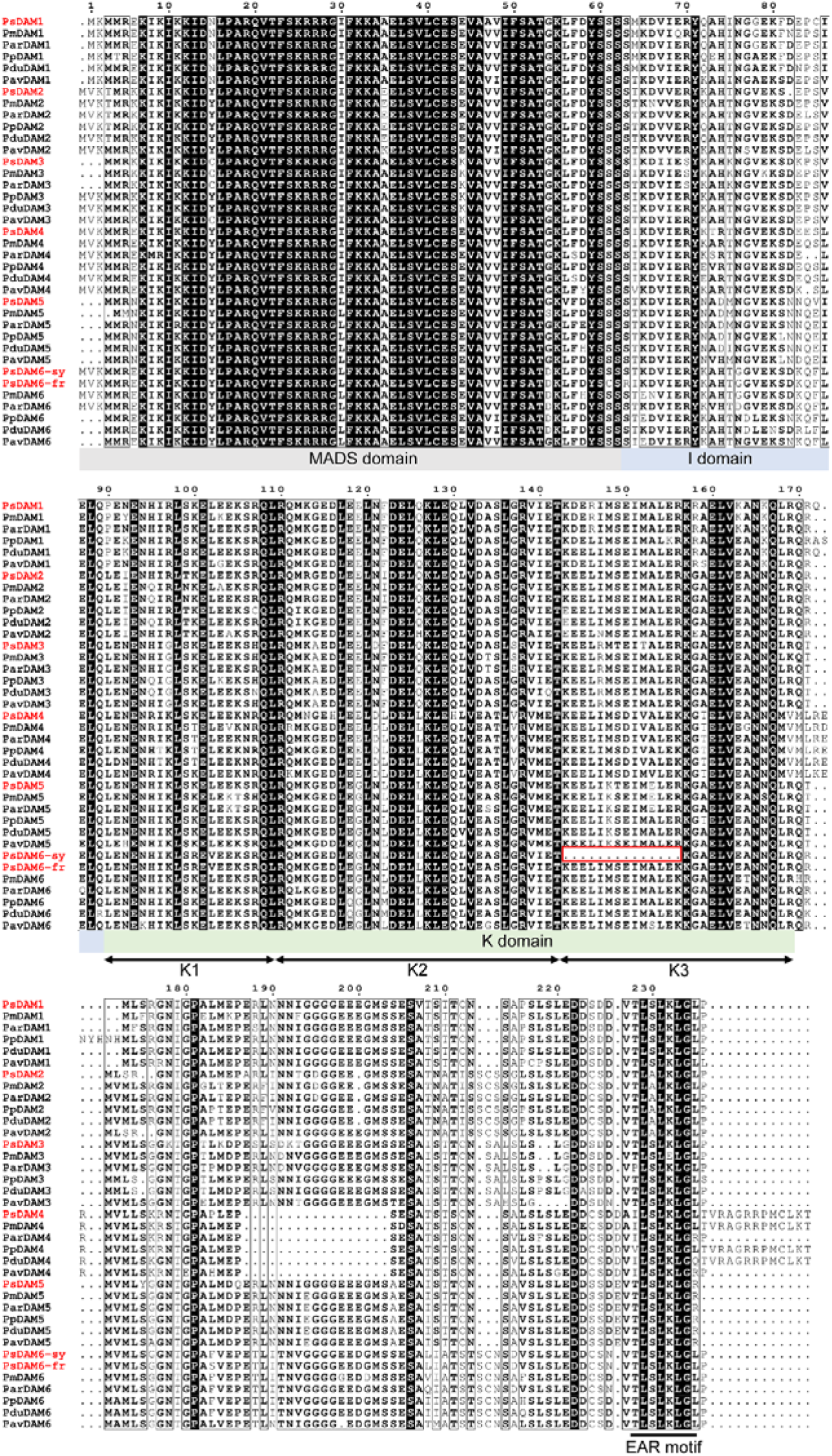
Multiple alignments of predicted amino acid sequences of *DAM* genes from plum and other *Prunus* species. Plum PsDAMs are indicated in red font. PsDAM6-sy and PsDAM6-fr represent PsDAM6 from ‘Sanyueli’ and ‘Furongli’ plum, respectively. The MADS, I and K domains are highlighted in grey, blue and green colors, respectively, at the bottom of the alignment. Three putative amphipathic a-helices, K1, K2, and K3, are indicated by arrows. EAR (ethylene-responsive element-binding factor-associated amphiphilic repression) motif is shown by lines on bottom of the alignment. Amino acids lost in PsDAM6-sy are indicated in a red box.

#### Transcriptome of dormancy release in plum flower bud

To investigate the molecular processes and genes involved in the regulation of flower bud dormancy, we compared the RNA-seq data derived from flower buds of ‘Sanyueli’ plum (low-chill) and ‘Furongli’ plum (high-chill) during chilling treatment. In total, 153.60Gb clean data were obtained and 85.4% to 92.0% of the clean reads from the libraries were successfully mapped to the genome of ‘Sanyueli’ (Supplementary Table 13).

Principal component analyses (PCA) showed that for each RNA-seq experiment, biological replicate samples tended to cluster together (Supplementary Fig. 7). Gene differential expression analysis identified 7782 DEGs with 3844 DEGs in F1 vs S1, 1313 DEGs in F1 vs F2, 1096 DEGs in F2 vs F3, 2213 DEGs in F1 vs F3, 1961 DEGs in S1 vs S2 and 1595 DEGs in S1 vs S3, respectively (Supplementary Table 14). However, only 288 DEGs were detected in S2 vs S3. KEGG enrichment analysis indicated that phenylpropanoid biosynthesis was one of the most enriched pathways in all comparisons except S2 vs S3 (Supplementary Fig.8). Starch and sucrose metabolism and plant hormone signal transduction pathways were also enriched in F2 vs F3 and F1 vs F3 (Supplementary Fig. 8).

The dramatic changes in the transcriptome of flower buds during chilling treatment involving a number of differentially-expressed genes indicate that the fulfillment of chilling requirement is a complex process of a series of transcription regulatory events. To identify candidate transcription factors that may participate in this process, we employed PlantTFDB v5.0 to predict transcription factors from the expressed transcripts. In total, 1,687 transcripts were predicted to encode transcription factors (Supplementary Table 15) and 511 of them were differentially expressed (Supplementary Table 16). All six *PsDAMs* were identified as DEGs (Fig. 6). *PsDAM1* and *PsDAM3* showed a similar expression pattern, with their expression only being significant in the buds of ‘Furongli’ at F3 stage and unchanged in the buds of ‘Sanyueli’. The expression profile of *PsDAM2*, *PsDAM4* and *PsDAM5* was similar in both cultivars during cold treatment. The expression of *PsDAM2* decreased in buds of both ‘Furongli’ and ‘Sanyueli’. The transcript levels of both *PsDAM4* and *PsDAM5* were increased by cold treatment and then decreased. However, the expression of *PsDAM4* was much higher than that of *PsDAM5*. The expression pattern of *PsDAM6* in flower buds of ‘Furongli’ during cold treatment was similar to that of *PsDAM4* and *PsDAM5*, but it is noteworthy that its expression in flower buds of ‘Sanyueli’ was much lower than that in flower buds of ‘Furongli’ and was significantly repressed after treatment with chilling temperatures for 170 h (Fig. 6). Thus, the expression of *PsDAM6* coincides with release of dormancy in flower buds of ‘Sanyueli’ and ‘Furongli’ plum.

**Fig. 6.**
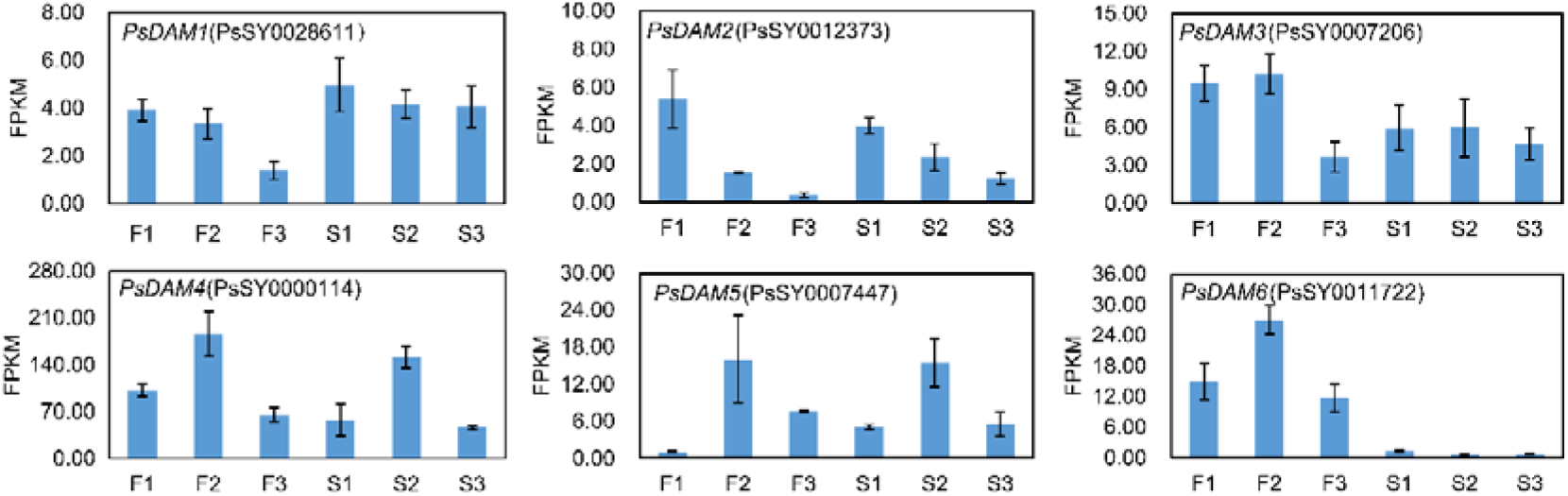
The expression profile of PsDAMs in dormant flower buds of ‘Sanyueli’ and ‘Furongli’ plum treated with chilling temperature. S and F represent ‘Sanyueli’ and ‘Furongli’, respectively. To search for genes that may regulate the expression of *PsDAM6* or regulated by *PsDAM6*, the correlation between the expression of *PsDAM6* and other differentially expressed genes was calculated. The expression of 190 differentially expressed genes showed a high significant correlation (|r| > 0.90) with that of *PsDAM6* (Supplementary Table 17). Some of them were lowly or not expressed in the flower buds of ‘Sanyueli’ (Fig. 7), including three transcription factors NAC (PsSY0001238), TT2-like (PsSY0001884) and LOB domain-containing protein (PsSY0011216). Interestingly, a transcriptional activator DEMETER-like protein encoding gene (PsSY0014977) was expressed well in flower buds of ‘Furongli’, but it was barely expressed in flower buds of ‘Sanyueli’.

**Fig. 7.**
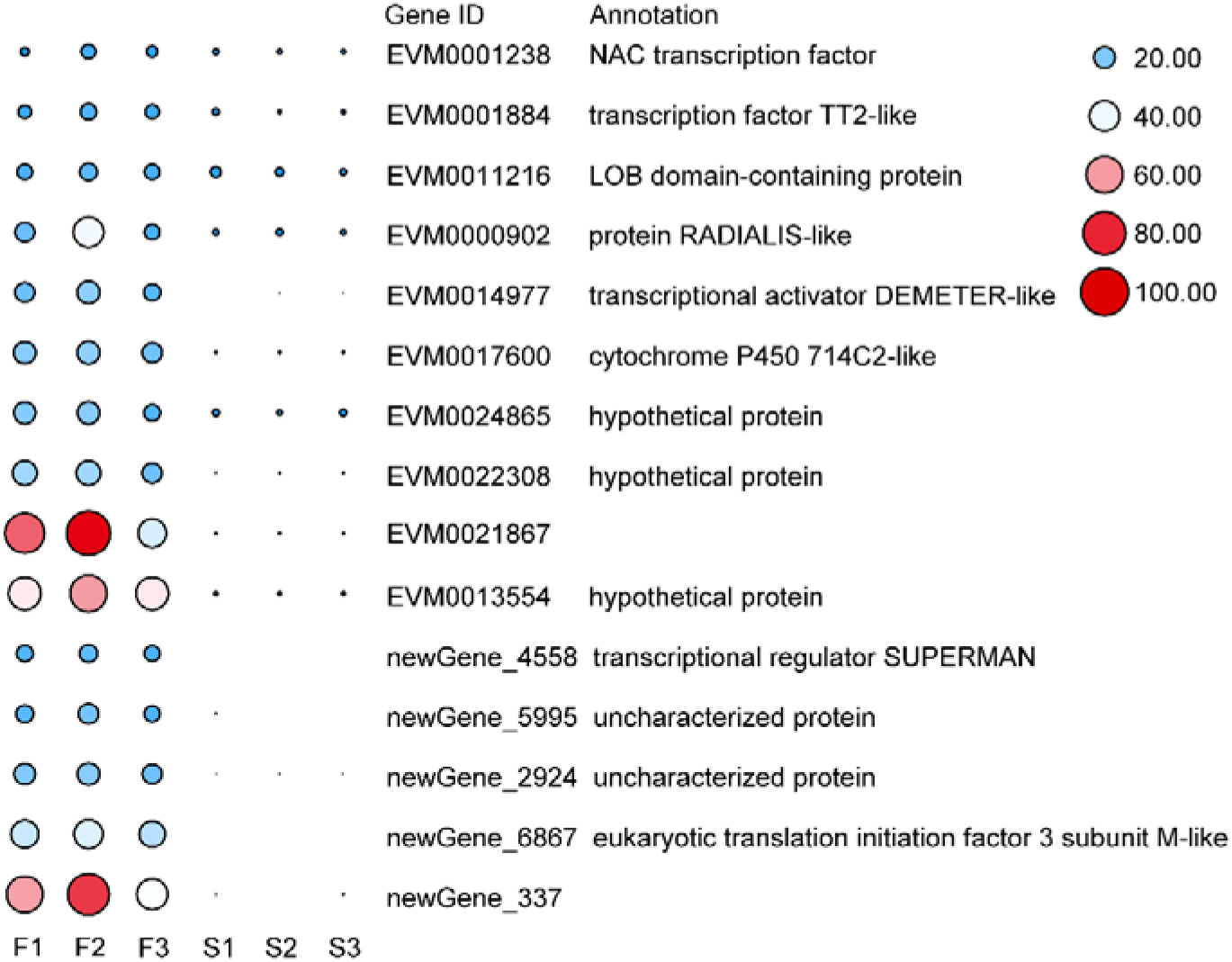
Expression profiles of selected differentially expressed genes correlated with PsDAM6. The expression levels of differentially expressed genes are shown as FPKM values.

## Discussion

Plums are widely grown for their fruits, which have excellent taste, nutritive value and processing compatibility^1^. Although Chinese plum is one of the most economically important stone fruits, its genome sequence has not been reported to date. In the present study, a 308.06 Mb genome of low-chill Chinese plum ‘Sanyueli’ with N50 of 815.7 kb was generated. The presence of the majority of BUSCO genes (97.0%) in our assambly indicated that the quality of ‘Sanyueli’ plum genome is comparable to that of sweet cherry (96.0%)^64^, almond(96.0%)^66^ and apricot(98.0%)^68^. The genome size of ‘Sanyueli’ plum is larger than that of mume^62^, peach^63^, almond^66^ and apricot (38.28%)^68^. 56.42% of the genome was predicted to be repetitive, which is much higher than that observed in other *Prunus* species, including flowering cherry (47.2%)^65^, mume (45.0%)^62^, sweet cherry (43.8%)^64^, almond (34.6%)^66^, apricot (38.3%)^68^ and peach (29.6%)^63^. The higher proportion of repetitive sequences (especially LTR insertions and expansions) may contribute to the larger genome size of ‘Sanyueli’ plum. 30,159 genes were predicted from the assembled genome of ‘Sanyueli’ plum, which is similar to that in apricot (30,436)^68^ and mume (31,390)^62^, but more than those in almond (27,969)^66^ and peach (27,852)^63^ and less than those in flowering cherry (41,294)^65^ and sweet cherry (43,349)^64^. The assembled ‘Sanyueli’ plum genome provides important information for understanding the evolution of *Prunus* species, the molecular basis of important agronomic traits and molecular assisted breeding of plum.

As global warming increases, unveiling the mechanism of flower bud dormancy and chilling requirements and the development of low-chill cultivars is of great importance for sustained production of plum. Low-chill cultivars of deciduous fruit trees have been used to identify genes related to bud domancy regulation^31,32,39,47,55,77,78^. Among the identified candidates, *DAM* genes were proposed as the central regulators controlling bud dormancy^14,33^. In the present study, *PsDAM* genes were found to be arranged in six tandem repeats in the genome of plum similar to *DAM* genes in peach and Japanese apricot^32,62^. Recently, six *PavDAM* genes were identifed in sweet cherry by Masuda, et al.^79^. Blast analysis results indicated there are six *DAM* genes, which were also arranged in six tandem repeats in the genome of *P. armeniaca* and *P. dulcis.* Sequence analysis showed that *DAM* genes from *Prunus* species shared similar gene structures and their predicted protein sequences were highly homologous to each other. These results suggested that *DAM* genes were conserved during the evolution of the *Prunus* family and that they may share similar functions. *PsDAM2*, *PsDAM4* and *PsDAM5* showed similar expression patterns in high-chill cultivar ‘Furongli’ and low-chill cultivar ‘Sanyueli’. Although the expression pattern of *PsDAM1* and *PsDAM3* was different between ‘Furongli’ and ‘Sanyueli’, their expression in ‘Sanyueli’ was not repressed by cold temperature. These results suggested that *PsDAM1-PsDAM5* are not responsible for the difference of chilling requirement between the cultivars. In peach, the expression of *DAM5* and *DAM6* is well correlated with the dormancy status of buds, highly expressed in dormant buds but lowly expressed in buds that have fulfilled their chilling requirements^33,80^. In addition, the expression of *PpDAM6* in buds of low-chill ‘Okinawa (Tsukuba)’ was lower and decreased earlier than that in high-chill ‘Akatsuki’^81^. Consistent with this result, our results demonstrated that the transcript level of *PsDAM6* was extremely low in buds of ‘Sanyueli’ and signifcantly decreased after 170h of cold treatment, while its expression in buds of ‘Furongli’ was approximately 10 times higher than that in buds of ‘Sanyueli’. Furthermore, *PsDAM6* was upregulated at 290h and decreased to a level comparable with that of the untreated control at 530h after cold treatment. Fan, et al.^31^ proposed that PpDAM5 and PpDAM6 could act as a dose-dependent inhibitor to peach bud break. Yamane, et al.^82^ indicated that break competency of dormant buds was significantly repressed in transgenic apple lines overexpressing *PmDAM6* from Japanese apricot. These results suggested that *PsDAM6* was the most significant candidate responsible for chilling requirement in plum.

Our results indicated that a large insertion in exon5 of *PsDAM6* in ‘Sanyueli’ resulted in partial deletion of the K domain, which is important for protein-protein interaction^83,84^. In Japanese apricot, Zhao, et al.^36^ demonstrated that PmDAM1 and PmDAM6 were shown to form heteromeric complexes with the cold response C-repeat binding factor PmCBF5. In addition, apricot PmuDAM6 was shown to interact with PmuSOC1^41^. Recently, similar interactions between PavDAM1/5 and the dormany-associated SUPPRESSOR OF OVEREXPRESSION OF CO1 PavSOC1 were reported by Wang, et al.^75^ in sweet cherry. The authors suggested these interactions between DAMs and SOCs could play a role in bud dormancy regulation. However, it remains unclear whether the lack of exon5 affects the function of the *PsDAM6* gene in ‘Sanyueli’.

Activation of *DAM* genes should be fine regulated in consideration of their role as dose-dependent inhibitors in the regulation of bud break^31^. Recently, transcription factors, such as peach PpTCP20 and pear PpyABF3 hve been shown to play roles in the control of bud dormancy by regulating the expression of *DAM* genes^85,86^. Our results showed that the transcript abundance of three genes predicted to encode transcription factors, NAC, TT2-like and LOB domain-containing protein, was positively correlated with that of *PsDAM6.* Conrad, et al.^58^ suggested that the phenylpropanoid pathway was associated with dormancy in apricot. Consistent with their results, transcriptome analysis results showed that DEGs were enriched in phenylpropanoid biosynthesis. TT2 is an R2R3 MYB activator involved in the regulation of proanthocyanidin accumulation in arabidopsis^87,88^. The differentially expressed TT2-like identified in this study could be responsible for transcriptional regulation of structural genes in the phenylpropanoid biosynthesis pathway. *NACs* have been demonstrated to be associated with dormancy release in seveval species, including peach^89^, pear^49,90^, grapre^25,91^ and peony^92–94^. Kumar, et al.^24^ reported that a NAC domain containing protein encoding gene was demethylated during the chilling acquisition. In addition Tuan, et al.^95^ indicated that NAC, as a cofactor of PpAREB1 repressed expression of *PpDAM1* in endodormancy release of pear buds. The differential expression of LOB domain-containing protein encoding genes has been reported during bud break of grape^25,96^. In sweet cherry, *PavLOB* was found to be highly expressed during endodormancy and around the time of dormancy release^53^. Epigenetic mechanisms were shown to participate in regulation of bud dormancy^55,97^. Several studies have reported that histone modification and DNA methylation are involved in transcriptional regulation of *DAM* genes during bud dormancy^28,48,51,76,97^. A gene predicted to encode DEMETER-like protein was found to be highly expressed in flower buds of ‘Furongli’ and expressed at an extremely low level in those of ‘Sanyueli’. Conde, et al.^98^ indicated that overexpression of DNA demethylase *CsDML*, a homolog of *PtaDML6*, significantly enhanced flavonoid accumulation through activating the expression of flavonoid biosynthesis genes and accelerated short◻day◻induced bud formation in poplar. Later, they demonstrated that a reduction of gDNA methylation in apex tissue during bud break is accompanied by the chilling-induced expression of *PtaDML10* and knock-down of *PtaDML8/10* delayed bud break in poplar. In addition, they further showed that the gene targets of DML-dependent DNA demethylation are genetically associated with bud break^99^. These results suggested that DML participated in the control of bud dormancy through modulating the methylation level of related genes. Further study will be required to determine wether the DEMETER-like gene is involved in regulating the expression of dormancy-related genes, such as *PsDAM6*, and flower bud dormancy in plum.

Several insertions were found in the introns of *PsDAM6* in low-chill plum ‘Sanyueli’. Falavigna, et al.^14^ proposed that intronic regions of *DAM* genes may function in the regulation of their transcription. Large insertions were observed in the first intron of both *PpDAM5* and *PpDAM6* in low-chill peach and was suggested to be linked to lower chilling requirements for dormancy release^81,100^. Saito, et al.^101^ reported a 3218 bp insertion in the first intron of *DAM* gene *MADS13-1*from low-chill pear ‘Hengshanli’. However, they found that the insertion also exists in a high-chill pear. Whether these insertions in the intron and exon region affect the expression and function of *PsDAM6* in flower buds of ‘Sanyueli’ requires further studies.

## Conclusions

In summary, we first report the sequencing, assembly and annotation of the genome of chinese plum ‘Sanyueli’, which is an extremely low-chilling requirement plum cultivar. Six PsDAM genes were identified in the plum genome. Transcriptome analysis suggested that *PsDAM6* was the key candidate responsible for the low-chilling requirement in plum. The genome of ‘Sanyueli’ plum provides a valuable resource for further research on the genetic basis of agronomic traits underpinning the genetic improvement of plum.

## Materials and methods

### Plant materials

Young leaves of ‘Sanyueli’ were collected from the plum repository at the Fruit Research Institute of Fujian Academy of Agricultural Sciences (Jinan District, Fuzhou, Fujian province, China). Annual branches of 8-year-old ‘Sanyueli’ were collected from the plum repository in Fruit Research Institute of Fujian Academy of Agricultural Sciences on December 7, 2017. Annual branches of 15-year-old ‘Furongli’ were collected from a commercial orchard in Gutian County, Fujian Province, China on December 5, 2017. The collected branches were immediately transported to the laboratory and placed in glass jars containing fresh water and kept at 2-6◻ in the dark. Flower buds of ‘Sanyueli’ were collected at 0 h (S1), 170 h (S2) and 450 h (S3) and flower buds of ‘Furongli’ at 0 h (F1), 290 h (F2) and 530 h (F3). To determine the dormancy state of the flower buds at each time point, a set of branches of ‘Sanyueli’ and ‘Furongli’ was kept at 25 ± 1°C with white light (150 mol m^−2^ s^−1^) under a 14 h light/10 h dark photoperiod at 75±5% humidity for 20 days. Three biological replicates for each sample were collected. Flower bud samples were immediately frozen in liquid nitrogen and then stored at −80°C until RNA extraction.

### Genome Sequencing

Genomic DNA was extracted from young leaves of ‘Sanyueli’ using a modified CTAB method. For short-read sequencing, a library with insert size of 270bp was constructed using TrueLib DNA Library Rapid Prep Kit for Illumina (Genetimes Technology Inc., Shanghai, China) and sequenced using Illumina HiSeq 4000 system (Illumina, San Diego, CA, USA). All raw reads were processed using a local perl script (developed by Biomarker Technologies Corporation) to remove adapter sequence and reads with length less than 100 bp or Q30<85%. For PacBio sequencing, genomic DNA was sheared into 20Kb fragments using a g-TUBE device (Covaris, Woburn, MA, USA), then the sheared DNA was purified and concentrated with AmpureXP beads (Agencourt, Beverly, MA, USA). A SMRTbell library was constructed with SMRTbell kits (Pacific Biosciences) according to the manufacturer’s protocol and sequenced on a PacBio Sequel system.

### Genome size estimation and heterozygosity

The filtered Illumina data were used to estimate the genome size, heterozygosity and the content of repetitive sequences through the k-mer depth frequency distribution analysis. This analysis was carried out using “kmer freq stat” software (developed by Biomarker Technologies Corporation, Beijing, China). Genome size G=K_num/Peak_depth, where the K_num is the total number of K-mer(k =19), and Peak_depth is the expected value of K-mer depth.

### Genome assembly

Raw PacBio reads were filtered using a local perl script (developed by Biomarker Technnologies Corporation). Only subreads equal to, or longer than 500bp were used for subsequent genome assembly. PacBio long reads were corrected and assembled using the Canu program^102^. In the correction step, Canu selects longer seed reads, then detects overlapping raw reads with MHAP (mhap-2.1.2), and finally performs an error correction through the falcon_sense method. Error-corrected reads were trimmed to obtain the longest supported range with the default parameters. Finally, a draft assembly was generated using the longest trimmed reads. In addition, we also used the WTDBG program (https://github.com/ruanjue/wtdbg) to assemble the Canu corrected reads. To improve the assembly contiguity, the assemblies generated by Canu and WTDBG were merged by Quickmerge (v0.2). Canu-generated contigs were used as query input and WTDBG -generated contigs were used as ref input. Finally, we employed Pilon (v1.22) to merge the assembly using high-quality cleaned Illumina reads.

### Evaluation of Genome Quality

To evaluate the coverage of the assembled plum genome, the Illumina paired-end reads were aligned to the genome assembly using the BWA-MEM (version 0.7.10-r789)^103^. To evaluate the completeness of the assembly, 1,440 Benchmarking Universal Single-Copy Orthologs (BUSCOs) and 458 Core Eukaryotic Genes (CEGs) were mapped to the plum genome using BUSCO v2.0^104^ and CEGMA v2.5^74^, respectively.

### Genome Annotation

The *de novo* repetitive sequence database of *P. salicina* was constructed employing LTR-FINDER v1.05^105^, MITE-Hunter^106^, RepeatScout v1.0.5^107^ and PILER-DF v1.0^108^. The database was classified using PASTEClassifier v1.0^109^ and merged with the Repbase database v20.01^110^ to create the final repeat library. Repeat sequences in *P. salicina* were finally identified and classified using RepeatMasker program v4.0.6^111^.

We used *Ab initio-based*, protein homology-based and RNA-seq-based approaches to predict protein-coding genes. *Ab initio-based* gene prediction was performed using Genscan v3.1^112^, Augustus v3.1^113^, GlimmerHMM v3.0.4^114^, GeneID v1.4^115^ and SNAP^116^; GeMoMa v1.3.1^117^ was employed to carry out homology-based prediction with the homologous peptides from *Oryza sativa*, *Prunus persica*, *Malus* × *domestica* and *Pyrus bretschneideri*. For RNA-seq-based prediction, the RNA-seq reads generated from leaf, stem and flower were assembled into unigenes using Hisat v2.0.4^118^ and Stringtie v1.2.3^118^. The unigenes were aligned to the assembly using BLAT^119^ and the gene structures of BLAT alignment results were modeled using PASA v2.0.2^120^. The prediction of protein-coding regions were performed with TransDecoder v3.0.1 (Haas, http://transdecoder.github.io) and GeneMarkS-Tv5.1^121^, respectively. Finally, the *de novo* predictions, protein alignments and transcripts data were integrated using EVM v1.1.1^122^. Annotations of the predicted genes were performed by blasting their sequences against a series of nucleotide and protein sequence databases, including KOG^123^, KEGG^124^, Pfam^125^, Swissprot^126^, NCBI-NT, NCBI-NR and TrEMBL^126^ with an E-value cutoff of 1e^−5^. Gene ontology (GO) for each gene were assigned by the Blast2GO^127^ based on NCBI databases.

The rRNA fragments were identified by aligning the rRNA template sequences (Pfam database v32.0) against the assembled plum genome using BLAST with E-value at 1e^−10^ and identity cutoff at 95% or more. The tRNAs were predicted using tRNAScan-SE v1.3.1 algorithms^128^ with default parameters. The miRNAs were predicted by INFERNAL v1.1 software^129,130^ against the Rfam database v14.0^131^ with a cutoff score of 30 or more. The minimum cutoff score was based on the settings, which yield a false positive rate of 30 bits.

### Comparative genomic analysis

Orthologous groups among *P. salicina* and 14 other species (including *Vitis vinifera*, *Rosa chinensis*, *Populus trichocarpa*, *P. persica*, *Prunus mume*, *Prunus dulcis*, *P. bretschneideri*, *Prunus avium*, *Prunus armeniaca*, *O. sativa*, *M.* × *domestica*, *Fragaria vesca*, *Amborella trichopoda* and *Arabidopsis thaliana*) were constructed using OrthoMCL v2.0.9^132^ based on an all-to-all BLASTP strategy (with an E-value of 1e-5). We extracted 436 single-copy genes from the 15 species and aligned proteins for each gene. All the alignments were combined into one supergene to construct a phylogenetic tree using RAxML v7.2.8^133^ with 1,000 rapid bootstraps followed by a search of the best-scoring maximum likelihood (ML) tree in one single run. Divergence time was estimated using the MCMCTree^134^ program in PAML v4.9 under the relaxed clock model. Several calibrated time points were used to date the divergence time.

A GF (gene family) was defined as a group of similar genes that descended from a single gene in the last common ancestor. Expansion and contraction of GFs were determined using CAFÉ v4.0^135^ based on changes in GF size. The cluster size of each branch was compared with the cluster size of the ancestral node. P value was calculated using the Viterbi method under the hidden Markov model, with p < 0.01 defining significant expansion or contraction. Genes belonging to expanded GFs were subjected to GO and KEGG enrichment.

All-against-all BLASTP analyses of protein sequences were performed between *P. salicina*, *P. mume* and *P. armeniaca* using an E-value cut-off of 1e^−10^. Syntenic regions between species were identified using MCScan^136^ based on the BLASTP results. A syntenic region was identified if it contained a minimum of 10 and a maximum of 25 genes in the identified gene pairs. Protein sequences of homologous gene pairs in the identified syntenic regions were aligned by MUSCLE v3.8.31^137^ and the protein alignments were then converted to coding sequence (CDS) alignments. The synonymous substitution (Ks) value of each syntenic gene pair was calculated using the Yn00 program in the PAML package^134^.

### Identification and analysis of *PsDAM*s

To identify *DAM* genes in the genome of ‘Sanyueli’, the deduced amino acid sequences of all six peach *DAM* genes^32^ and six *P. mume PmDAM* genes^61^ were used as queries for Blast analyses using TBtools^138^. The schematic gene structure of *DAM* genes and structural alignments of the *PsDAM6* gene from ‘Sanyueli’ and ‘Furongli’ plum was displayed using TBtools^138^. Phylogenetic analyses were performed using the neighbor-joining method by MEGA-X software with 1,000 bootstrap replicates. The predicted amino acid sequences of *DAM* genes in other Rosaceae species were downloaded from Genbank or Genome Database for Rosaceae, including *M. domestica* MdDAM1 (KT582786), MdDAM2 (KT582787), MdDAM3 (MDP0000527190), MdDAM4 (KT582789), MdJa(KT582788), MdJb (LC004730); *P. bretschneideri* Rehd. PbrDAM1 (KP164027), PbrDAM2 (KP164026) and PbrDAM3 (KP164028); *P. persica* PpDAM1 (DQ863253), PpDAM2 (DQ863255), PpDAM3 (DQ863256), PpDAM4 (DQ863250), PpDAM5 (DQ863251) and PpDAM5 (DQ863252); *P. armeniaca* ParDAM1 (PARG08688m02), ParDAM2 (PARG08688m01), ParDAM3 (PARG08689m02), ParDAM4 (PARG08689m04), ParDAM5 (PARG08690m03) and ParDAM6 (PARG08690m02); *P. avium* PavDAM1(LC544139), PavDAM2 (LC544140), PavDAM3 (LC544141), PavDAM4 (LC544142), PavDAM5 (LC544143) and PavDAM6 (LC544144); and *P. mume* PmDAM1 (XR_001677199.1), PmDAM2 (XM_008221048.2), PmDAM3 (XM_008221049.2), PmDAM4 (XM_008221050.2), PmDAM5 (NM_001293268.1) and PmDAM6 (NM_001293262.1). DAM genes were predicted from the genome sequence of *P. dulcis* in the Genome Database for Rosaceae according to the sequence of peach *DAM* genes, PduDAM1 (Pd01:39769437-39776175), PduDAM2 (Pd01:39778066-39783736), PduDAM3 (Pd01:39795512-39804580), PduDAM4 (Pd01:39805806-39814186), PduDAM5 (Pd01:39815116-39823593) and PduDAM6 (Pd01:39824719-39831140). Alignments of DAM proteins were generated with ClustalW and displayed using ESPript 3.0^139^.

### Transcriptome sequencing and analysis of flower buds

Total RNA was extracted from approximately100 mg flower buds using the RNAprep Pure Plant Kit (Tiangen, Beijing, China). The quality of the RNA was assessed using an Agilent 2100 Bioanalyzer (Agilent Technologies, Santa Clara, CA, USA). The concentration of RNA was determined using Qubit 2.0 Fluorometer (Invitrogen, Life Technologies, CA, USA). High quality RNA was subjected to library preparation using NEBNext Ultra™ RNA Library Prep Kit for Illumina (NEB, USA). The libraries were sequenced on a HiSeq X Ten sequencer (Illumina). The RNAseq reads have been deposited into the NCBI Short Read Archive and are accessible under project PRJNA645255. The RNA-seq reads were mapped to the ‘Sanyueli’ plum genome sequence using HISAT2^140^. Reads were assembled into transcripts and the expression level of transcripts was calculated as the FPKM values using the StringTie software package^141^. Principal component analyses (PCA) were performed on FPKM values from different datasets using the GenAlEx version 6.5 program. Differential gene expression analysis was carried out using DESeq2^142^. Genes with fold change greater than 2 and a false discovery rate (FDR) below 0.05 were considered significantly differentially expressed. GO enrichment analysis of differentially expressed genes (DEGs) was performed with the GOseq R package. GO terms with corrected p-values < 0.05 were considered to be significantly enriched by differentially expressed genes^143^. KOBAS software was used to test the statistical enrichment of DEGs in KEGG pathways^144^.

The PlantTFDB v5.0 (http://planttfdb.cbi.pku.edu.cn/prediction.php) was used to predicted transcription factors from all assembled transcripts. The correlation coefficient between the expression of *PsDAM6* and differentially expressed genes in buds of ‘Sanyueli’ and ‘Furongli’ plum was calculated using Microsoft^®^ Excel^®^ software.

## Supporting information

Supplementary Fig. 1 Daily maximum temperature and minimum temperature from November 2015 to February 2016

Supplementary Fig. 2 Bud break of cuttings of plum trees after treated with chilling temperature

Supplementary Fig. 3 Distribution frequency of the 19-kmer graph for genome size estimation

Supplementary Fig. 4 Distribution map of genes from the three prediction methods

Supplementary Fig. 5 Collinear analysis of P. alicina, P. armeniaca and P. mume genome

Supplementary Fig. 7 Principal component analyses of the RNA-Seq samples

Supplementary Fig. 8 KEGG pathway enrichment analysis of the annotated DEGs

## Acknowledgements

We thank Rongmei Wu for critical reading of the manuscript. This research was funded by the National Natural Science Foundation of China (31801916), the Basic Scientific Research Funds of Public Welfare Scientific Research Institutes of Fujian Province (2018R1013-1), the China Scholarship Council (CSC) (201909350001) and Projects of Fujian Academy of Agricultural Sciences(YC2015-9, STIT2017-1-4 and AGY2018-3).

## Data availability

The raw genomic sequence and transcriptome data have been deposited in the NCBI Sequence Read Archive under accession numbers PRJNA650137 and PRJNA645255

## Contributions

Z. F. and X. Y. designed and managed the project. H. D. performed assembly and annotation of plum genome; Z. F. analyzed the RNA-seq data. Z. F., D. Z., C. J., Y.L. and S.P. prepared and handled samples. Z. F. drafted the paper. Y.X., K.L., and R.E. revised the paper. All authors read and approved the final version of the manuscript.

---

**Supplementary Fig. 1 Daily maximum temperature and minimum temperature from November 2015 to February 2016.**

**Supplementary Fig. 2 Bud break of cuttings of plum trees after treated with chilling temperature.** A,‘Sanyueli’. B, ‘Furongli’. The pictures were taken at 20 days after being transferred to and kept at 25 ± 1°C with white light (150 mol m^−2^ s^−1^) under a 14 h light/10 h dark photoperiod at 75% humidity.

**Supplementary Fig. 3 Distribution frequency of the 19-kmer graph for genome size estimation.** Density plot of the frequency of unique 19-kmer for each kmer depth (x axis) is plotted.

**Supplementary Fig. 4 Distribution map of genes from the three prediction methods**

**Supplementary Fig. 5 Collinear analysis of P. salicina, P. armeniaca and P. mume genome**

**Supplementary Fig. 6 Alignment of genomic DNA sequences of the *PsDAM6* from ‘Sanyueli’ and ‘Furongli’ plum.** Exons are highlighted in green shading and different bases between the *PsDAM6* from ‘Sanyueli’ and ‘Furongli’ plum are indicated in red font. The genomic DNA sequence of *PsDAM6* in ‘Furongli’ plum was extracted from the genome of ‘Furongli’.

**Supplementary Fig. 7 Principal component analyses of the RNA-Seq samples.**

**Supplementary Fig. 8 KEGG pathway enrichment analysis of the annotated DEGs.**

