## Supplementary figures and images for "The genome of low-chill Chinese plum ‘Sanyueli’ (*Prunus salicina* Lindl.) provides insights into the regulation of the chilling requirement of flower buds"

### Supplementary Fig. 1 Daily maximum temperature and minimum temperature from November 2015 to February 2016

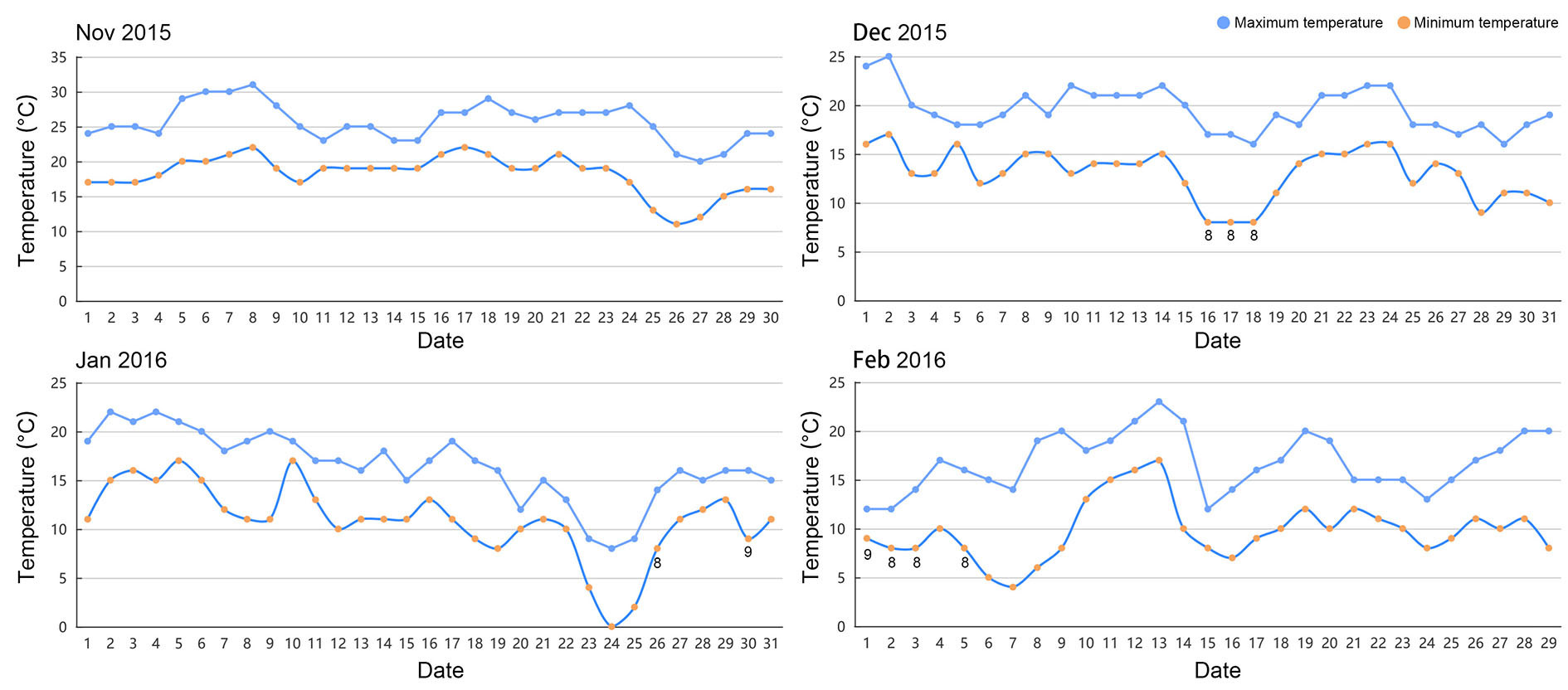

### Supplementary Fig. 2 Bud break of cuttings of plum trees after treated with chilling temperature

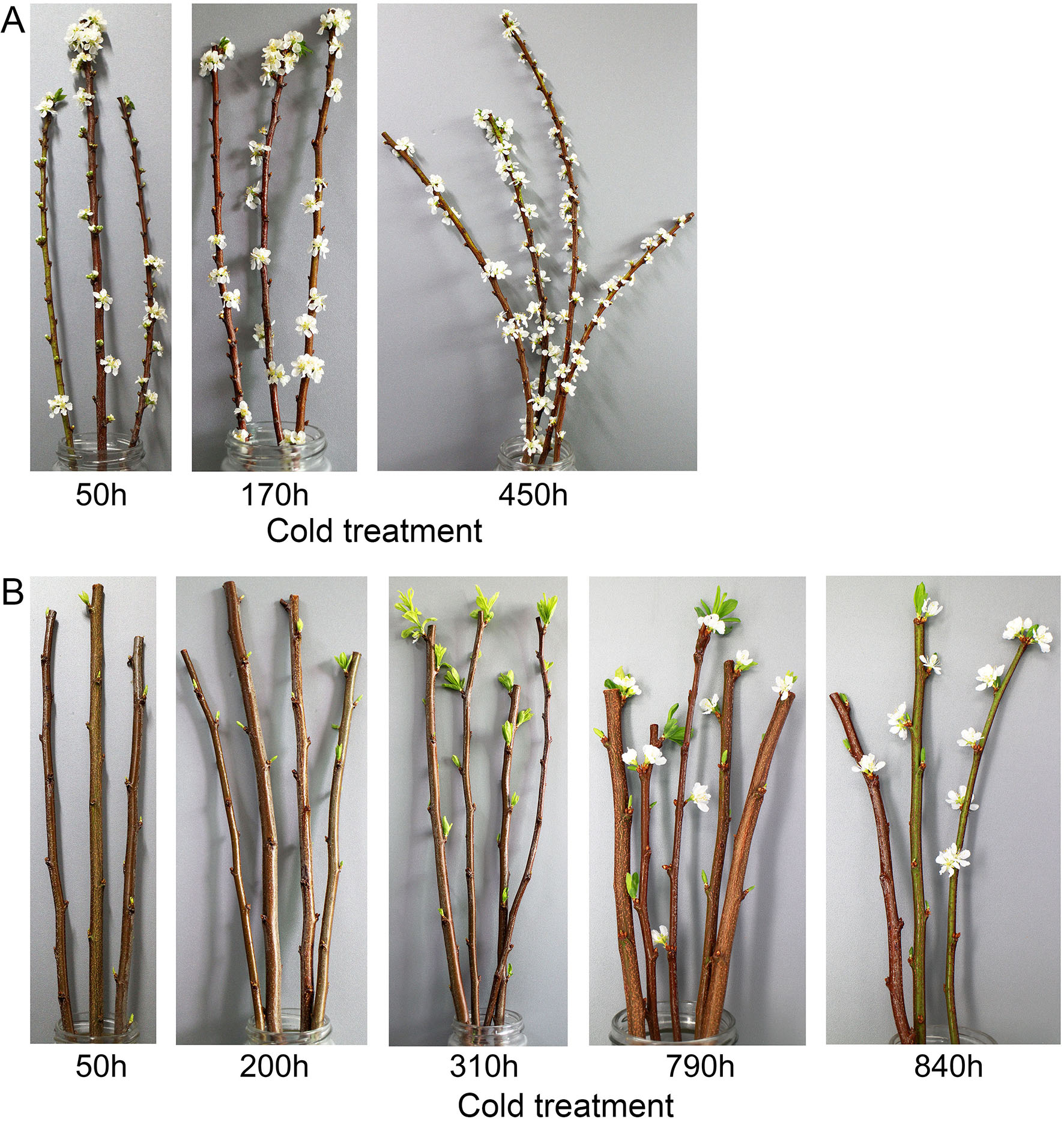

### Supplementary Fig. 3 Distribution frequency of the 19-kmer graph for genome size estimation

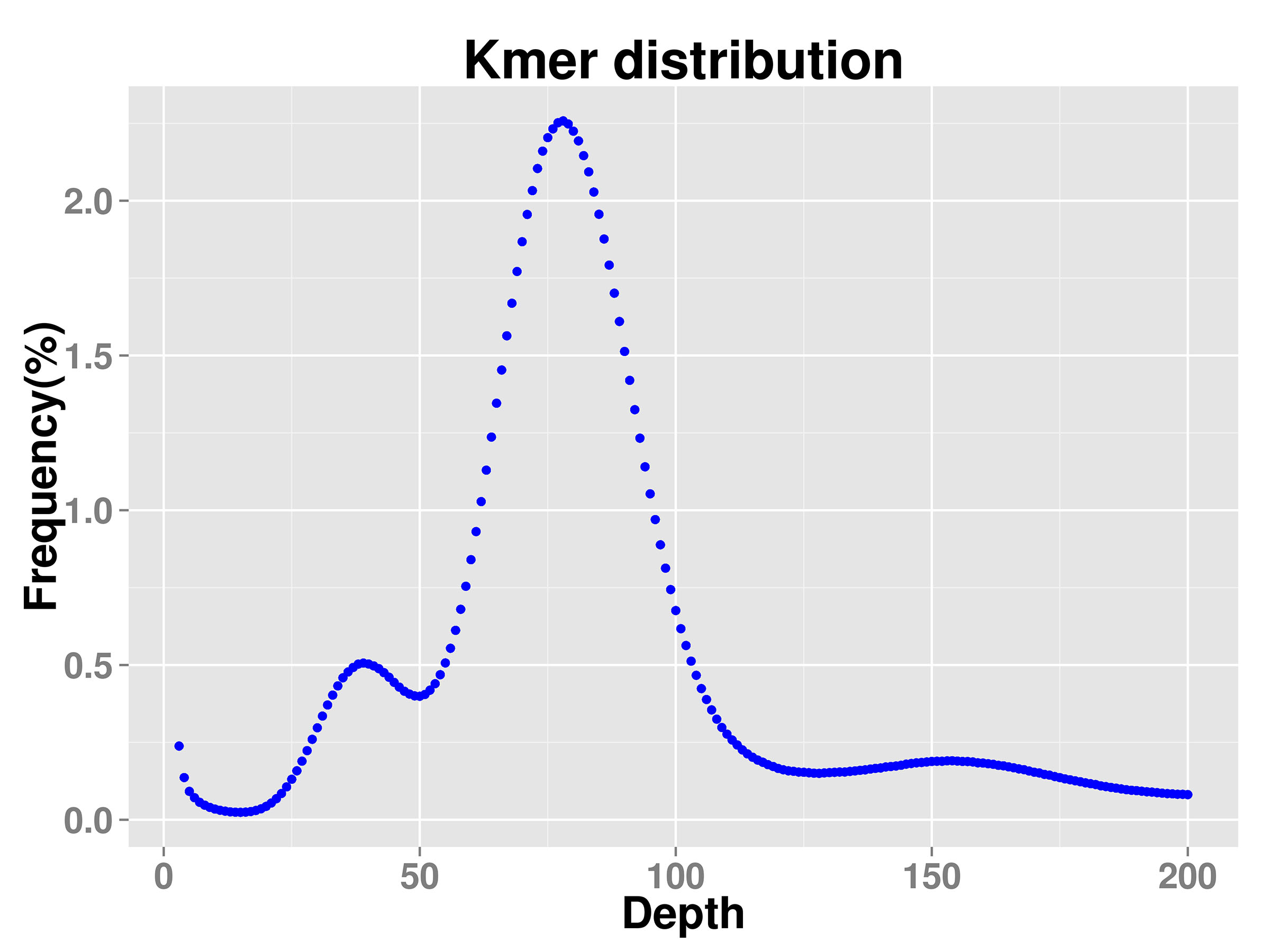

### Supplementary Fig. 4 Distribution map of genes from the three prediction methods

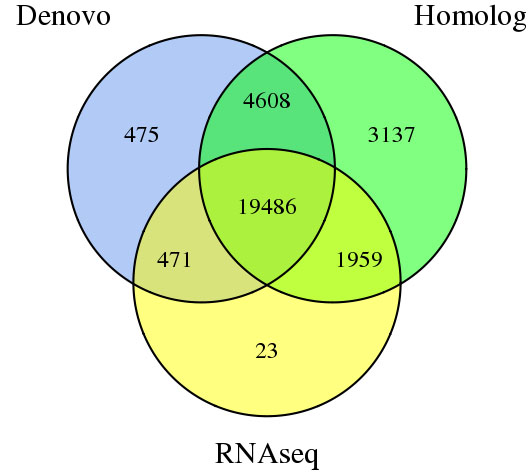

### Supplementary Fig. 5 Collinear analysis of P. alicina, P. armeniaca and P. mume genome

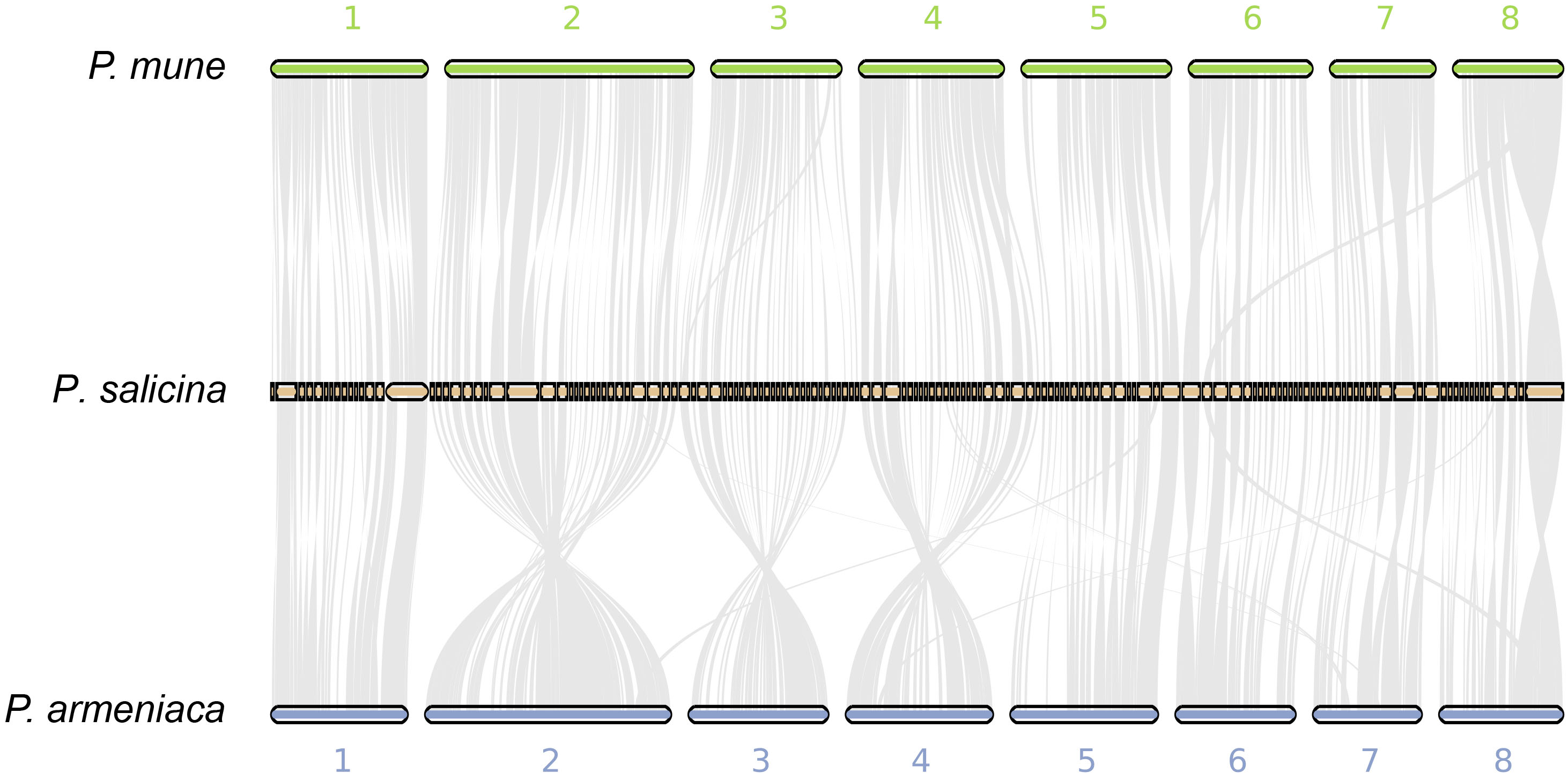

### Supplementary Fig. 7 Principal component analyses of the RNA-Seq samples

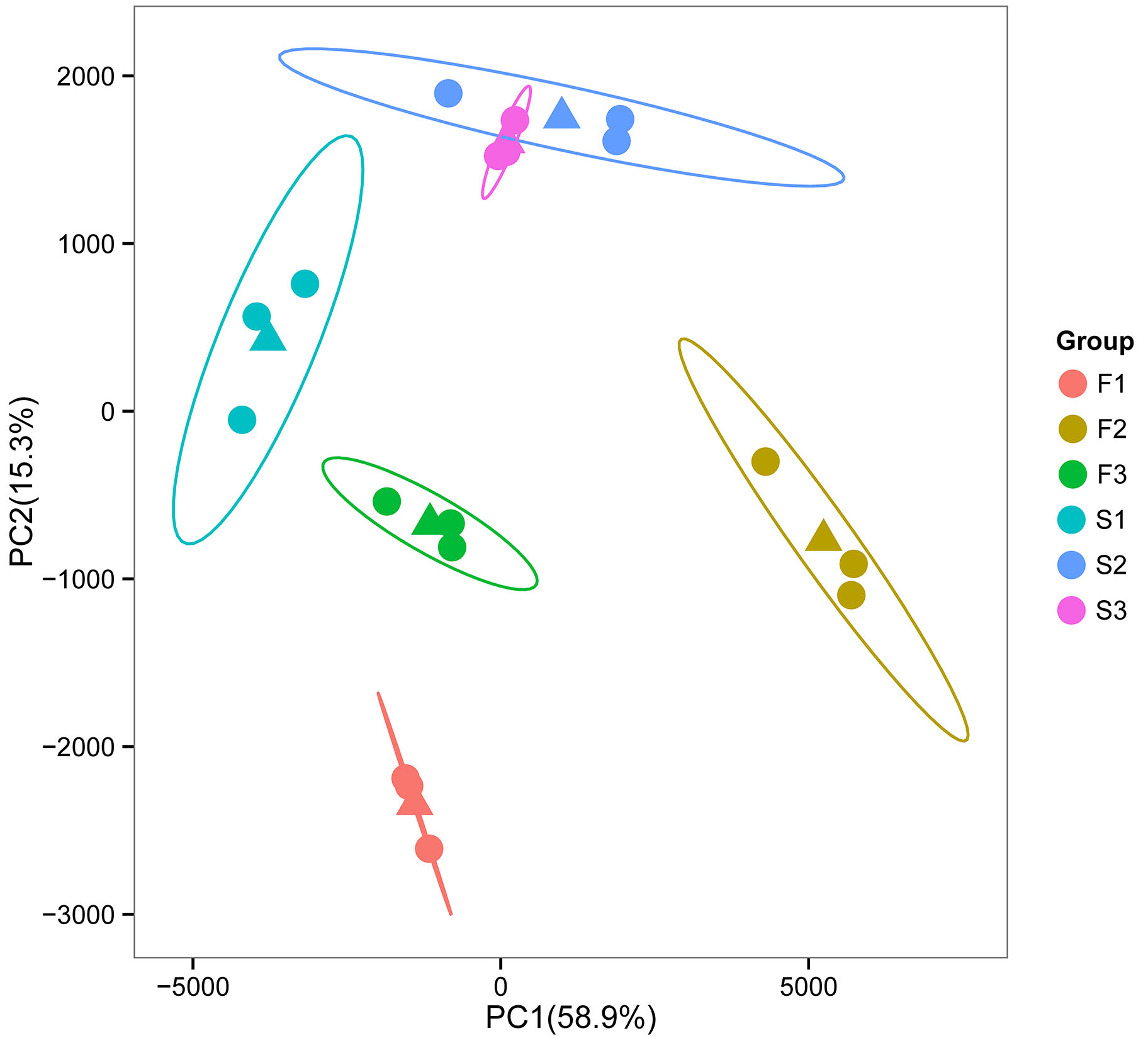

### Supplementary Fig. 8 KEGG pathway enrichment analysis of the annotated DEGs

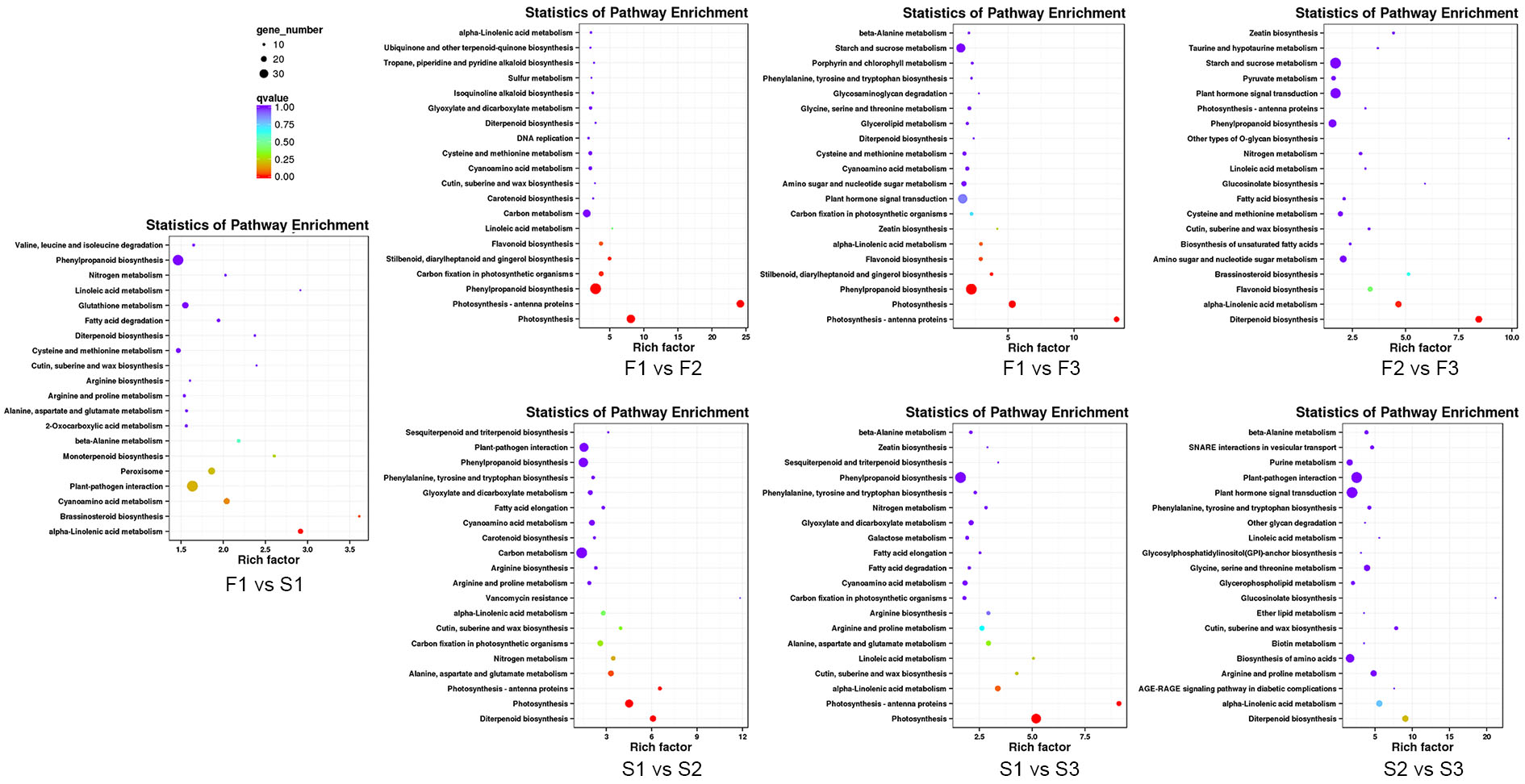
