## Supplemental Data 1 for "The genome of low-chill Chinese plum ‘Sanyueli’ (*Prunus salicina* Lindl.) provides insights into the regulation of the chilling requirement of flower buds"

```
SY    ATGGTGAAAATGATGAGGGAGAAGATCAAGATCAAGAAGATTGACTACCTGCCAGCAAGGCAGGTTACCTTTTCAAAGAGAAGAAGAGGGCTCTTCAAGAAAGCTGCAGAGCTATCTGTT
FR    ATGGTGAAAATGATGAGGGAGAAGATCAAGATCAAGAAGATTGACTACCTGCCAGCAAGGCAGGTTACCTTTTCAAAGAGAAGAAGAGGGCTCTTCAAGAAAGCTGCAGAGCTATCTGTT

SY    CTGTGTGAATCTGAGGTGGCTGTCGTCATCTTTTCTGCTACTGACAAGCTTTTTGATTATTCAAGCTCAAGGTACCAGCACCCACCACTATATACCTTATAATTCTTTTGACTCTTTCCT
FR    CTGTGTGAATCTGAGGTGGCTGTCGTCATCTTTTCTGCTACTGACAAGCTTTTTGATTATTCATGCTCAAGGTACCAGCACCCACCACTATATACCTTATAATTCTTTTGACTCTTTCCT

SY    TCCTTGAAAAGATAACCAATATGTATGATCATGATGTCTGAAGTTTGACTTGACTTGTATGTGTGTGT----------------------GAGAGAGAGAGAGAGAGAGAGAGAGAGAGA
FR    TCCTTGAAAAGATAACCAATATGTATGATCATGATGTCTGAAGTTTGACTTGACTTGTATGTGTGTGTGTGTGTGTGTGTGTGTGTGTGTGAGAGAGAGAGAGAGAGAGAGAGACTGAGA

SY    GAGAGAGAGACTGAGATTCAGTCATTTGGATAGAAATATATTTAGAGGAGAGGGTGCCTTTTATGGAAATTACCTGTCCTTCCTGAAATTCCATGGTTTAAACTCTGCACACAGACATTA
FR    ----------------TTCAGTCATTTGGGTAGAAATATATTTAGAGGAGAGGGTGCCTTTTATGGAAATTACCTGTCCTTCCTGAAATTCCATGGTTTAAACTCTGCACACAGACATTA

SY    TTTATTGTCTCTAAGCTTCCAAAAGATAAAAAGGAGAACCAGAAGCTTCTAGAGAGAGAGGGAGAGAGAGAATCTTG------ATAACTTTTAGATAGTTATACTCTATTTCAAAGGCAA
FR    TTTATTGTCTCTAAGCTTCCAAAAGAAAAAAAGGAGAACCAGAAGCTTCTAGACAGAGAGGGAGAGAGAGAATCTTGTAGATGATAACTTTTAGATAGTTATACTCTATTTCAAAGGCAA

SY    CTAGTTCGAGGGTTACTACCTTGTATACGTTTGATTAAGACAACTTCTAGCTAGGTGTCGCAGTTTGAGAGACATTTATATGGTGATGTTCTTTTCTTGTACTACATGCGCAATCAATTT
FR    CTAGTTCGAGGGTTACTACCTTGTATACGTTTGATTAAGACAACTTCTAGCTAGGTGTCGCAGTTTGAGAGACATTTATATGGTGATGTTCTTTTCTTGTACTACATGCGCAATCAATTT

SY    TGTTTACCAAGCAAAACTGTGTATATCTTTTTGGACAAGTGTATATATAAAATCTGTAATCTTATCCTCCACCTATCACATATTGGTAGTAGATATTATATAAAATCTTAGGTGGGGCCT
FR    TGTTTACCAAGCAAAACTGTGTATATCTTTTTGGACAAGTGTATATATAAAATCTGTAATCTTATCCTCCACCTATCACATATTGGTAGTAGATATTATATAAAATCTTCGGTGGGGCCT

SY    ATACAACGAAGAATATCTTTGTTCCAAAAGGACATAAAATTAATGGTTTTCTTGTGAGAGACACGGTATATTAGACCATCAATTAAATCATAAACTCAAACAAGAATGTAAATCACCAAA
FR    ATACAATGAAGAATATCTTTGTTCCAAAAGGACATAAAATTAATGGTTTCCTTGTGAGAGACACGGTATATTAGACTATCAATTAAATCATACACTCAAACAAGAATGTAAATCA-----

SY    TCTAGATTTGTAGTTTGACCGCATAAAATGATCGATAGTCTAGCTAGTAAGAGAGTATTTGACTCTTTCAGATATATACATATCTATACTATATGTATATATCTTATCCTTCAAACTTAA
FR    ---------------T-ACCGCATAAAATGATCGATAGTCTAGCTAGTAAGAGAGTATTTGACTCTTTCAGATATATACATATCTATACTATATGTATATATCTTATCCTTCAAACTTAA

SY    GATAATGTTGGGTTGTGTTTGAAAAGGCGTCTTTCACTCTCCCCGCCTAAACTAAAAAAATGCCAGTAAATTAATGTGGTTACAAACATTGAATTATATGTTTGTAGTTTTCTTACTAAA
FR    GATAATGTTGGGTTGTGTTTGAAAAGGCGTCTTTCACTCTCCCCGCCTAAACTAAAAAAATGCCAGTAAATTAATGTGGTTACAAACATTGAATTATATGTTTGTAGTTTTCTTACTAGA

SY    AATAATGTGATATAATGATAACTTGAGTATTCATCATATAGAAGCCCTGAAGACTTTCAGCAGGAATATGCATGCATGGTTTCAAGTAGGGCAATGGGTGGTGGAGCTATGACCAT----
FR    AATAATGTGATATAATGATAACTTGAGTATTCATCATATAGAAGCCCTGAAGACTTTCAGCAGGAATATGCATGCATGGTTTCAAGTAGGGCAATGGGTGGTGGAGCTATGACCATAACA

SY    --------CCAT-AA-----------------------CATGTTTCATGCTTAGATGTGCATACATGAGAGGGAGGGTGAGAATGAAATGTTAACTATGGAATAAGAATTTAAATAAGGT
FR    TGTTTTTGCCATGAATGCATGAAAGATCTTGTTAAAATCTAGTTTCATGCTTAGATGTGCATACATGAGAGGGAGGGTGAGAATGAAATGTTAACTATGGAATAAGAATTTAAATAAGGT

SY    CATGCTTCCTTTCAGTTAATACCACCTCTGTTTTTTAGAAAAAAGAATTCCGTTATTTAGATGTTCCTATTTTAGTATTAA---------------------------------------
FR    CATGCTTCCTTTCAGTTAATACCACCTCTGTTTTTTAGAAAAGAGAATTCCATTATTTAGATGTTCCTATTTTAGTATTAACTAGCCTCTCTGCATGCGCTTCCGCGCCTGCAAGAGGCC

SY  --------------------------------------------------------------------------------------------------------------------------
FR    TTTTTTAAAAGAATTTAATTTTTTTTTTAAGAATTAAAAAGATAATGGATAGTTGTGTTCCGTAAAAATAGGATCCATTATCTTTTTCTTTTAATTTTAATTTTTTTTAGAAGTGTGAAT

SY    ----------------------------------------------------------------------------------------------------------------------TA
FR    TTACCATATTATTATCATTTAATTAATAATTTTAATTCTAATATTTGAATTAACCAAGGGCATTTTTTGGTATTTTGAATGTTTCACCATTCTCTACCTTTTGCTTTATATATATAGATA

SY    ACATGCGAAAAATGAAAGAAACAGATTAAAAGACAATTTGAAAATGCAAATGCGAATTAAGAACAAAATTAAAAGACCTTTTAACTGTTTATGTCAGTATGCAATTTCTAATCAATTTCC
FR    ACATGCGTCAAATGAAAGAAACAGATTAAAAGACAATTTGAAAATGCAAATGCGAATTAAGAACAAAATTAAAAGACCTTTTAACTGTTTATGTCAGTATGCAATTTCTAATCAATTTCC

SY    ATTGAAAATTTCATGTAAAATTTATTTTTCTAGGTTTCCGGCATAATTTGACACGTTATATGCATGAAAAGCTGCTAGTTTGTCTAGTTTGTCTA-TTCTGTATTTTTTTTTTTTCAAGT
FR    ATTGAAAATTTCATGTAAAATTTATTTTTCTAGGTTTCCGGCATAATTTGACACGTTATATGCATGCAAAGCTGCTAGTTTGTCTAGTTTGTCTAGTT-TGTATTTTTTTTTCTTCAAGT

SY    TTTTCTATTAAAGCTCTCGATATTACAGTCCATGTTTTCATTTTTGCAACCAAAACAAAACAAAAAAAACATAAAACGATTTTACTCAAATCAATATACATGGAAAATCTAGGGCATGTT
FR    TTTTCTATTAAAGCTCTCGATATTACAGTCCATGTTTTCATTTTTGCAACCAAAACAAAACAAAAAAAACATAAAACGATTTTACTCAAATCAATATACATTGAAAATCTAG--------

SY    TACGTATTAGTAATGGGAATGAGAGGAAAGGGAATGATTTCCATTCCTGACTTCTCCTGCGTTTACTTGCAATCAGGAATGAAAAAATAAGTGGGCCCCATCTCAAAATCCGTAATCACA
FR    ------------------------------------------------------------------------------------------------------------------------

SY    TTCCCTCAAAAGTAGGAATCAGATCAACTAGGGGGGAGTAGTCGATACGATTCCCATTCCTGTTTTCTCTTAATAACTTCCAAAAATACCCTTATCTCTTTACCATAATTCTAAAATTAA
FR    ------------------------------------------------------------------------------------------------------------------------

SY    TTTGGATTCTGATTCACGCAAATTAGTAAACAATAACAATGGGAATCAATCTCATTCCATTGCTAATTCCATATGTTTAGTAAACAGCTTAATGGGAATGATTCTTCCCATACCGATTCA
FR    ------------------------------------------------------------------------------------------------------------------------

SY    GTAGTTCTCCGATTCCTAAATTCTCTGATTCCTTAGTTTCTCCATTACTGATACGTAAATGTGCCCCTAGAGGAGAATAAAAGTGTTGTTCTTTAGCCTTAGATTTTACTTTACATTATT
FR    ----------------------------------------------------------------------AGGAGAATAAAAGTGTTGTTCTTTAGCCTTAGATTTTACTTTACATTATT

SY    TTTCAAATTTTTCCAAGTTTCCAAGTAGAGCAATATTGGTCCAAATTTTCCAATTATAAATAGGGCTTGTTAAGCGAGGCCATGTTAATCCAAAACATGATTCTAAGTTTTCTATAAGGG
FR    TTTCAAACTTTTCCAAGTTTCCAAGTAGAGCAATATTGGTCCAAATTTTCCAATTATAAATAGGGCTTGTTAAGCGAGGCCATGTTAATCCAAAACATGATTCTAAGTTTTCTATAAGGG

SY    TGGGTGCAGCCCGATTCTCACCCCAAATTGGAATGAGAATTTGAACTATTCATCTAGTTCGATTCAATTTGGTTTTGACCAAAAAAATTCTCCGGTCTAGTGTGGTATGATTTGCCCATG
FR    TGGGTGCAGTCCGATTCTCACCCCAAATTGGAATGAGAATTTGAACTATTCATCTAGTTCGATTCAATTTGGTTTTGACCAAAAAAATTCTCTGGTCTAGTGTGGTATGATGTGCCCATG

SY    AGTAGGATCGATTTACGATTTTGGATCGGAAATTTTTTTTTTGTACAAAACTCAATTAAAATTTTATTTTTTGGGCTTAAAAATTCACAAATGGCCCAATTTCATGCAAAAGAAGATGTG
FR    AGTAGGATCGATTTACGATTTTGGATCGGAAATTTTTTTTT-GTACAAAACTCAATTAAAATTTTATTTTTTGGGCTTAAAAATTCACAAATGGCCCAATTTCATGCAAAAGAAGATGTG

SY    ATTGGGCCTAGAAAAGAGAGGTGATTTGTAGTTGGGCTTGGAGAAAGATTAATTATGGGCTTAGAAATTTTGCATTGGGTTAAATTTTTTATTAAAAACTTAAATAATAAATAGTCGGTT
FR    ATTGGGCCTAGAAAAGAGAGGTGATTTGTAGTTGGGCTTGGAGAAAGATTAATTATGGGCTTAGAAATTTTGCATTGGGTTAAATTTTTTATTAAAAACTTAAATAATAAATAGTTGGTT

SY    CGAATTGGCTCAATCCGATTTTTTCACTACAAGAATCAAAATCAAAATCAAACCAAACCAGTACGGTTTGGTTTTTAGTCAACTCGGTTTTTTTTGGCCTAAATCAGCTCTGTTTATGCC
FR    CGATTTGGCTCAATCCGATTTTTTCACTACAAGAATCAAAATCAAAATCAAACCAAACCAGTACGGTTTGGTTTTTAGTCAACTCGGTTTTTTTAGGCCCAAATCAGCTCTGTTTATGCC

SY    CACCACTAGTTTTCTACTCGATTTTCATGTCCTATTAGCCTAGCTTTGGGGAGTCTAGATCCCCTTGTTGCAGTAGGGCTTGATTAAGTTATTCTTCAAGAATCAATGGTTTAAAAGGGT
FR    CACCACTAGTTTTCTACTCGATTTTCATGTCCTATTAGCCTAGCTTTGGGGAGTCTAGATCCCCTTGTTGCAGTAGGGCTTGATTAAGTTATTCTTCAAGAATCAATGGTTTAAAAGGGT

SY    CTATTAATATATCTACTAAGGATTCGGCCCCCAAGTGATGTGTGACGTTTCCGACGTGTTTAATGATCTTTCTAGGGTGCATCTGGCTTCTGAAGCTCTAAAGTGTTTGAGAAGAACAAA
FR    CTATTAATATATCTACTAAGGATTCGGCCTCCAAGTGATGTGTGACGTTTCCGACGTGTTTAATGATCTTTCTAGGGTGCATCTGGCTTCTGAAGCTCTAAAGTGTTTGAGAAGAACAAA

SY    ATTAGTTAAGAATGAAATGTTATAGCCTCTAAGGTGTGTTAACTGCATTTTAGTTCTTCGGTGTCTATGGCCGTACTGAGGAATGGAGGAGATCACACGCCCGGACATAGGAGAAAGATG
FR    ATTAGTTAAGAATGAAATGTTATAG--TCTAAGGTGTGTTAACTGCATTTTAGTTCTTCGGTGTCTATGGCCGTATTGAGGAATGGAGGAGATCACACGCCCGGACATAGGAGAAAGATG

SY    ACGCTGGGATGATGGTCATGGATTTATTCACCTAGGTAGAGACTTTTGATATTTTTGGCGCAAGCCAGTAGAAGTTTCAACTTGATATTGCAACATAAAATTAATAACAATATTCACTTA
FR    ACGCTGGGATGATGGTCATGGATTTATTCACCTAGGTAGAGACTTTTGATATTTTTGGCGCAAGCCAGTAGAAGTTTCAACTTGATATTGCAACATAAAATTAATAACAGTATTCACTTA

SY    TTTGTGATATCGAAGGTGTATGAATATGTCATTCGGATTTAGGAAACAACTCTTAGAGAAAAC-TATT-AATATTC--------------------------------------------
FR    TTTGTGATATCGAAGGTGTATGAATATGTCATTCGGATTTAGGAAACAACTCTTAGAGAAATTGTGTTTTCTTTTGAAGATACAGGATTTAGGAAACAACTCTTAGAGAAATTGTGTTTT

SY    -----GAAGATACATAAGTGAACAATCAATCTATTAATATTCGAAGGACAATGTTTTTCCAATATATTGTTTGTCAATCTTTTCTACTA----------ACCCCCCAACCCAAGGAAAAC
FR    CTTTTGAAGATACATAAGTGAACAATCAATCTATTAATATTCGAAGGACAATGTTTTTCCAATATATTGTTTGTCAATCTTTTCTACTAAGATCGAATGACCCCCCAACCCAAGGAAAAC

SY    TGAATTCGAATTCAGGTACATCTTTTAGGGCGAAATCTTTTAGGTACATC-TTTTCTACTATA------TATACTTGTATTTAGCCACCTTCCACATGATAGCCTTTAGAAAAAGGATTG
FR    TGAATTCGAATTCAGGTACATCTTTTAGGGCGC------------ACATCTTTTTTCACTAAAAGTGGCTATACTTGTATTTAGCCACCTTCCACATGATAGCCTTTAGAAAAAGGATTG

SY    CGAAATCTCAAATATCTCTCGATGCTGAGAATTTTCTATGTGTCCAGGATTTCTATTAGTGGAGATACCTATTTTTATACGGCTGGATAAAGCTTTTTCGAATCTTCAATCCATTAGTAT
FR    CGAAATCTCAAATATCTCTCGATGCTGAGAATTTTCTATGTGTCCAGGATTTCTATTAGTGGAGATACCTATTTTTATACGGCTGGATAAAGCTTTTTCGAATCTTCAATCCATTAGTAT

SY    AAGAAAAACGGTGGACTCTCCCAAGAGCATATGAGTGCGCAAGATCTAATATATCAACAACTGATTCTACTTAGATGCTTTTATCCGCTTCACACTTGGCTACCCATCGTTTATAGAAGT
FR    AAGAAAAACGGTGGACTCTCCCAAGAGCATATGAGTGCGCAAGATCTAATATATCAACAACTGATTCTACTTA--TG-------------------------------------------

SY    AGCTTACTCATCTAGCGGCAAAGGAAAAGGGTAGTACTTTGAGTTATTTTATTAGGCGGGGACATGATCCGAACCTATGCCTGTCACAAAGAACTTAGGAAAGTATCTTAGCGTGCCACT
FR    ------------------------------------------------------------------------------------------------------------------------

SY    AATTCACATGCAAAAGATACACCATTAGAGTAAGTGCACTTTACTAACATAAAATAAAGAAAGTTGGAGTTTCAGGATTTAGGGTTTTAGTTCAGGAATTAAGGTTTAGGGTTTAGGTTT
FR    ------------------------------------------------------------------------------------------------------------------------

SY    AGGCTGAAGACATGGGTTTTGGAGAGAACTACTAAAGGTGTGTATAAGCCCTGCACACATCGTATGTTATCCTAAAGGTTCATAATGGATGGGGGTTGAGGTTAAAATCTAAGACCTACA
FR    ------------------------------------------------------------------------------------------------------------------------

SY    TATGACCTAACCCGAAGCGGATAGTTTTTTTTTTTCTCAATTATATTTTGCACTCATGTGTCTAGAGTTTGTGGTAGTCTATCCCTAACTGCTATAGGGAGAGATGAGAATAGCCCAAGT
FR    ------------------------------------------------------------------------------------------------------------------------

SY    GCCTTCAATTTTGCATTCATGTAAACGAAAAAAGGGGTTAGTTTGACCGTTAGATGGGCGCAATTGTGTAACAGTTTGTAATGTTATGATTGTTTTTTGCTAGTTTTTAAAACACATTAT
FR    ------------------------------------------------------------------------------------------------------------------------

SY    CGGTAGTATTATTGGCATAGGCCTACATAGAGTTGTGTGTGTAGATAGTCGAAATAAAAAGAGAGCGAGAATTGTCTCACTACCCTGCGAGAAAAACTTGGGTCCGATTGATTTTCATGA
FR    ------------------------------------------------------------------------------------------------------------------------

SY    ATTAATTTAGTGCTTGAGATTCGTATTTGCATGGTAGCTAGCGCTGACGAATTCAGGATAAAAAGAAGCCAATCAAAAACACCCACTGACTAAAAAGAGGCCAACAAAGATAACAATATT
FR    ------------------------------------------------------------------------------------------------------------------------

SY    CAAAGCATATTTCAATGAGCTCCAGAGTTGATGTGAGAGGAACTGTCTATAGCTAAGATTTTTGAGGCCCACTTATTAGCTGACTTGATTTTCTCACGTCGGCTTGACTTGTGTTCTAAA
FR    ------------------------------------------------------------------------------------------------------------------------

SY    ATATCTCAGCTGCTTGGGTTGGTCTTATATATAATAATTCCTTAAAATAGCTATGAAGGAATTAGAGTTGACGTTTTGATTTGCATATGTAACCAAACCGAGCAATAAGGCCAGCAGCAA
FR    ------------------------------------------------------------------------------------------------------------------------

SY    ATAGCTTAACGTTCTCATTATGCATATCAAATAAAAGTATATGTATAAGTGTATATCTGTGTTTGCAAACCAGGAAGTTTGAATAATAGGGATAAATTCTTTTCCATTTTTTTGTTTCTC
FR    ------------------------------------------------------------------------------------------------------------------------

SY    ATGTATGTTTAGTCACACGTTAATTTTTCTATGTTTAATAATTTGCTAGTGAACATGCCAATTGATTGTTTTTTTTACCTTTTATGTTGGGAAGTACCAAGGATGTTATTGAAAGGTACA
FR    ------------------------------------------------------------------------------------TGT-----AGAATCAAGGATGTTATTGAAAGGTACA

SY   AAGCGCACACAGGTGGTGTTGAAAAATCGGACAAACAGTTTCTTGAGCTGCAAGTAAATGAGTTCAATGACCTTTTCTCTTTCCTCTTTTAAATTGTTCTGCAATTAATTGTAACATTACT
FR   AAGCGCACACAGGTGGTGTTGAAAAATCGGACAAACAGTTTCTTGAGCTGCAAGTAAATGAGTTCAATGACCTTTTCTCTTTCCTCTTTTAAATTGTTCTGCAATTAATTGTAACATTACT

SY    CTGTGATCTGCACTTTGGAGATTCAAAATGTATCTCTATAGCAACTGACGGTTCTGATTGTTTGAACATAAAATAAAAAAACATTATCCTGTTACATGCCAGACTTATGTGTATGCATGC
FR    CTGTGATCTGCACTTTGGAGATTCAAAATGTATCTCTATAGCAACTGACGGTTCTGATTGTTTGAACATAAAATAAAAAAACATTATCCTGTTACATGCCAGACTTATGTGTATGCATGC
FR    ************************************************************************************************************************

SY    AGCAGGGTCGGCATCTTAATTGGGATATATATATATATATATATGTAGGTGGCTTCGATAAGTCGATTTCTAGTGGACTTGATTTGGTCTACATAAGAATTAGCCAATGAAAAAATTGGG
FR    AGCAGGGTCGGCATCTTAATTGGGATATATATATATATATATATGTAGGTGGCTTCGATAAGTCGATTTCTAGTGGACTTGATTTGGTCTACATAAGAATTAGCCAATGAAAAAATTGGG

SY    ACATAGATTCATCATCAACTATCAATTATTATTTACTTTTACTGTTAGTTGTTTTCCAAGTGTTCAAAGGAGAGGGTAAATTATTTTATAGGTTTTTACAAATAAGTCCATTATATAGAG
FR    ACATAGATTCATCATCAACTATCAATTATTATTTACTTTTACTGTTAGTTGTTTTCCAAGTGTTCAAAGGAGAGGGTAAATTATTTTATAGGTTTTTACAAATAAGTCCATTATATAGAG

SY    GCTATTTGAAGATAAAAACTCAAAATTAGCCACTTGTCATTTTCTGATTGGTTTTTAGAAAATTTGAACACACTAATAACCAATCAGAAAATGACAAGTGGCTAGTTTTAGGTTTTTCTC
FR    GCTATTTGAAGATAAAAACTCAAAATTAGCCACTTGTCATTTTCTGATTGGTTCTTAGAAAATTTGAACACACTAATAACCAATCAGAAAATGACAAGTGGCTAGTTTTGGGTTTTTCTC

SY    TCCAAATAGCCTCTATATGCGTTTTTGTATAGTATGTCATGTTAGAATGGACATATTTGTGAAAAGTCTACTATTGAATTGTTTTTCAAGCTTTAGGGGATTTAATTTCTTATGATTTTT
FR    TCCAAATAGCCTCTATATGCGTTTTTGTATAGTATGTCATGTTAGAATGGACATATTTGTGAAAAGTCTACTATTGAATTGTTTTTCAAGCTTTAGGGGATTTAATTTCTTATGATTTTT

SY    TCAAAACTCCAGATTGTTGAGTTGGAGACTTCTATGTGTCCCTCGCACTTTGGTTGCAGTTGAGGCTTAGCCCCTAAATAACCTCTTTCACAGGCTCACATAGTACCCTAATTTTGTTAG
FR    TCAAAACTCCAGATTGTTGAGTTGGAGACTTCTATGTGTCCCTCGCACTTTGGTTGCAGTTGAGGCTTAGCCCCTAAATAACCTCTTTCACAGGCTCACATAGTACCCTAATTTTGTTAG

SY    CTACCTAAATCCTGACAAAACTAGGAAATCAATAAATACCCATAACGACTAGGACAAATTGTAAATCCATATTCTTGTATGTTCCAGTAATTCATTTGATTCCATTTTCAAAAATATATT
FR    CTACCTAAATCCTGACAAAACTAGGAAATCAATAAATACCCATAACGACTAGGACAAATTGTAAATCCATATTCTTGTATGTTCCAGTAATTCATTTGATTCCATTTTCAAAAATATATT

SY    GTAATCCTTTTCGTTCTTCTCGTTATGTTGTTTCTTTGTTTTTTGTTTTTTGTTTTTTGTTTTTTGTTTTTTTTGGGGGGCTGATGTTTATAAGCTTTTTATGTGACAGATTAGATTATG
FR    GTAATCCTTTTCGTTCCTCTCGTTATGTTGTTTCTTTGTTTTTTTT-----------------------------GGGGGCTGATGTTTATAAGCTTTTTATGTGACAGATTAGATTATG

SY    TAATGGTTCCAATATTTCTCATTTTAGCTGGAGAATGAAAACCACATCAAACTGAGTAGGGAAGTCGAGGAGAAGAGCCGCCAGCTGAGGTACAAATTTATTAAATCATCATGAACTTCT
FR    TAATGGTTCCAATATTTCTCATTTTAGCTGGAGAATGAAAACCACATCAAACTGAGTAGGGAAGTCGAGGAGAAGAGCCGCCAGCTGAGGTACAAATTTATTAAATCATCATGAACTTCT

SY    AAGATATATGTGGCTGTTTAGCTGGTATATTATTGCTTGAATTATGTGCGGGAATTGGACTGGCTTGCAACAGGCAGATGAAAGGTGAGGATCTTGAAGGGCTGAATCTTGATGAGCTGC
FR    AAGATATATGTGGCTGTTTAGCTGGTATATTATTGCTTGAATTATGTGCGGGAATTGGACTGGCTTGCAACAGGCAGATGAAAGGTGAGGATCTTGAAGGGCTGAATCTTGATGAGCTGC

SY    TGAAGTTAGAACAACTGGTGGAAGCAAGCCTTGGCCGTGTGATAGAAACTAAGGTCTGCCCTAGCTAATTTCTTTTCTAGTGAGGATGGTGCAAGGCGTCACTCTCTGTATTGTGGTTTA
FR    TGAAGTTAGAACAACTGGTGGAAGCAAGCCTTGGCCGTGTGATAGAAACTAAGGTCTGCCCTAGCTAATTTCTTTTCTAGTGAGGATGGTGCAAGGCGTCACTCTCTGTATTGTGGTTTA

SY    ATGATCGAAACCGTTTATTATAATTTAGGAATCCTTGTTGCAGAATAGTTAAATGGTGTTAGTACAACTGACAGTCTAAATTATAGGGGAGAGAGGTTTCTCACACACACTCACACGACT
FR    ATGATCGAAACCGTTTATTATAATTTAGGAATCCTTGTTGCAGAATAGTTAAATGGTGTTAGTACAAGTGATAGTTTAAATTATAGGGGAGAGAGGTTTCTCACACACACTCACACGACT

SY    AAGATGCCAATGGGGTTTCGAATCCGAGACCCCTAGTCTGCAAATTAAAGTTATTTTCCACTGGGTTAGACCC-GTTAGCTTAGTT-AAATGGTGCAAGACATTGCATTTGTGTTCCAAT
FR    AAGATGCCAATGGGGTTTCGAACCCGAGACCCCTAGTCTGCAAATTAAAGTTATTTTCCACTAGGTTAGACCCCATTAGCTTAGTTTAAATGGTGCAAGACATTGCATTTGTGTTCCAAT

SY    AAATAAACAA--GCATCTCAACAAAACACTGAAATCATTACCATTTAATGAAATGGTCAAATAGTATCCAATATTCTATTTTTTTGCCGGATGATTCTTAAATGAAAACCTAAATGATGA
FR    AAATAAACAACAGCATCTCAACAAAACACTGAAATCATTACCATTTAATGAAATGGTCAAATAGTATCCAATATTCTATTTTTTTGCCGGATGATTCTTAAATGAAAACCTAAATGATGA

SY    ACTGTTCGAGTAAAGTAGGTCCCACAATTGTCCTTTGTTTGAAGAAGTTGTTTGTGCATAGAGAGTATATTTTAATTGCTCTATTTTGTGAAAAGCATAAGTGTATCCTGGGTTTTCACT
FR    ACTGTTCGAGTAAAGTAGGTCCCACAATTGTCCTTTGTTTGAAGAAGTTGTTTGTGCATAGAGAGTATATTTTAATTGCTCTATTTTGTGAAAAGCATAAGTGTATCCTGGGTTTTCACT

SY    AAGGTTTGAGTTGCCAATTTGGGTTACATTATGCAAATAAACTAACATCAAAATGCAAATGTTCCATTCCTTTTCAGGAAGAGCTGATTTTTTTTTTTTTTTTTTTTTTTTTTTTTTTTG
FR    AAGGTTTGAGTTGCCAATTTGGGTTACATTATGCAAATAAACTAACATCAAAATGCAAATGTTCCATTCCTTTTCAGGAAGAGCTGATT-------------------------------

SY    GTCAATGGAAGCTTTCAATCAAGAAACGAAACAAGACAACAAAGGGGAGAAACAAACAAACCAAAGACAGAAAGAAAAGGAACAAGCAAACAACAGCAAAAAAACACACAGCAGCTAGAA
FR    ------------------------------------------------------------------------------------------------------------------------

SY    AACTAAAAGCTAGAGACCAGTAGCTATGACGTCTCCAAAAGGGGAGCCCGTCCGCCTCCAGCACACACTCCCTAGCACCGCGGGGGGCAAGGAAGTCCGTCAGCTTGCAACACATAAACC
FR    ------------------------------------------------------------------------------------------------------------------------

SY    AGAGAGGATGGGGGGCGGGAAGCCCAAACTTCATCGCACATCCTCCTTCTAGCTCGCTGAGCAGCAGCATCCGCTGCTCTGTTTGCCCCTCGAGGGACCCAGTTCCAAGAACAAGAACTA
FR    ------------------------------------------------------------------------------------------------------------------------

SY    AAACTGCTACAAAGTCGCCGAATAGCTGATAGGATAGGATAAATCATCCACCTACCTTTCAAAACATTTCCTTTAACTGATTGAACGAGTTCCTTGGAGTCAGATTCAAAGACCACGTTT
FR    -----------------------------------------------------------------------------------------------------------------------

SY    GGAAACCCCCTTTCAGCAGCCAGCTTAACCCCCTTTAAAGCAGCATGGGCTTCAGCCTCAATTGCAGAAGAAGCAAGCCTAGTGGCGCAGCAACCACCCATAAAGTTACCAGCAGAATTT
FR    ------------------------------------------------------------------------------------------------------------------------

SY    CTCACGACAACTCCCAGACCTGTTTGCAAAGTACCACTAACCCAAGAAGCATCAATATTTATTTTGTAAACATAAGTGGGAGGGGGGGACCATCTAACCACCCTATGGCCATCATCAATA
FR    ------------------------------------------------------------------------------------------------------------------------

SY    TTCTCCTCCAAAATCACATCCCCAGGAGAACCCCACACAAGATTGAAATCATTTATCAAATTCTTTGCCACCAGCAAAGTACGACGAGGGCACACACTAATATCATCAAATAAAGCAGAG
FR    ------------------------------------------------------------------------------------------------------------------------

SY    CAACGGCTTTTCCAGATACTCCAACAAGTGTAAACAATTTGGGAAAGTAGCCACTTTCGGTCATATCCCACACCCTGCGAAAACTTCAAAATCTGCATCATCCAATCACTCATAGAAGTA
FR    ------------------------------------------------------------------------------------------------------------------------

SY    ATACTCTGTCTATTTATCCTGTAGTTCAAAGGACCTCCAAACCATACAGGCTGGACCCAATTACAGAGGAGGAAAAGATGCTCAACTGACTCCGGAAACTCCCCACAAATAGGGCATAAC
FR    ------------------------------------------------------------------------------------------------------------------------

SY    GGGGAGGTGCCAAGGTGTCTCCTGAACAAAGCATCCCTTGTTGGCAGGCAGCCCCTGACCAATCTCCACCAAAAATTCATTAACTTTGGGACCATCTGAGAGCCCCAAATCAGCTTCCAT
FR    ------------------------------------------------------------------------------------------------------------------------

SY    ATAGCCTTGTCTAAAACTCTTGAGCTGGATGGCCTGACCGGAGTACCAAGGTGAGCCATGTGAATCATGTTATAGCCAGATTTGACAGTATAACTCCCCGATTTGTTTAGAGGCCATATA
FR    ------------------------------------------------------------------------------------------------------------------------

SY    AGCCTGTCCTTCTCCCAACCATCACCCAAAGGCATGGCTCTGATAATTTTTGAGGCATTTGGTGAAAACATACCACCAATAGCCTCCAAATTCCATTCACGAGATTGACAATTAATAATA
FR    ------------------------------------------------------------------------------------------------------------------------

SY    GTCTCAACCTTTGCCTCCAAATCCACCTGGCTGAGCTGGCTTGACTGCAGAGCATGCTCAGGACACCCGGGGATCCACTTATCAGTCCAAAGGTGAACACGGCTACCATCAAGAACCTGC
FR    ------------------------------------------------------------------------------------------------------------------------

SY    CACCTAGCACCTCTCATGATTATATTCCTTCCTACCAGCAGGCTAGACCAAGCCCATGACGACTTAGCCCCCTTCCTGGCCCCGAGAAAATCACCATTCGGGAAATATTTACTTTTTAGT
FR    ------------------------------------------------------------------------------------------------------------------------

SY    AGTTGAGCCCAAAAAGCCTGAGGCTCCATCACCATCCTCCACCCTTGTTTTGCCAACAAGGCAACATTAAACTCCTTCAGGTTGCGGAAGCCCATTCCCCCTTCATTCTTAGGCATGCCA
FR    ------------------------------------------------------------------------------------------------------------------------

SY    AGGTCCTTCCAACTAATCCAATGAATTTTATTTGATTGCTGGGATTGTCCCCACCAGAAATTTGCCAAAATACTATCAATCTCCTGACAGAAGCCATTCGGGAAGAGGAAGACAGACATA
FR    ------------------------------------------------------------------------------------------------------------------------

SY    GGGTAAGACGGGACGGCCTGGGCAACAGACTTAATCAGGACCTCCCTGCCAGCTTGAGAGAGTAAGCCATGCTTCCAACCCTGGATTTTTCCCAGAATTTTATCCTTTACAAAAGCCAAA
FR    ------------------------------------------------------------------------------------------------------------------------

SY    GCCATCTTTTTTGATCTGCCCCATAAAGTTGGAAGCCCAAGATACTTGCCCGGATCCTCAGAGATGGTAACATTAAGAATAGCCCTCAACCTATCCTTCACCTCCAAAGGCGTATTTGGG
FR    ------------------------------------------------------------------------------------------------------------------------

SY    CTGAAGAACATATTTGATTTCTCAAAATTAACAAGCTGACCCGAGGCCGTGCAATAAGATTCCAAAATTCTCACAATCACCCGACAGTTATTCTCAGATGCTTTTAGAAACATAAGTGAG
FR    ------------------------------------------------------------------------------------------------------------------------

SY    TCATCAGCAAAAAACAGGTGTGGATTGATCATGGAAGAGAGCACATCACTAACGATAATAAACAAGTATGGAGATAAAGGATCCCCTTGTCTGAGCCCTCTTGTCGGAGTGAAATAGCTT
FR    ------------------------------------------------------------------------------------------------------------------------

SY    CCCACTTTCCCATTGACAAGTACAGCAAATTCAACAGAAGATAAGCACCTCATGACCCATCCTACCCATTGGCAAGCAAAGCCCATCTTTAACAGCACCGCCTGAACAAAATCCCACTCG
FR    ------------------------------------------------------------------------------------------------------------------------

SY    ATCCGATCGTAAGCCTTACTCATGTCCAGCTTGAGCCCCATCTCAAAAATTTTTGTTTTCCTCCTAATCTTTAAGGAGTGGAAAGCTTCATGAGCCACCAAAACATTATCTTGGATCTGT
FR    ------------------------------------------------------------------------------------------------------------------------

SY    CTCCCTGGAATGAAAGCACATTGTTGAGGCGAGATGAATTTATCCAAGAATGGCTGCAGTCGATTTGCTAAAATCTTTGAAATAATCTTATAGGAATAATTACAAAGGGAGATAGGACGG
FR    ------------------------------------------------------------------------------------------------------------------------

SY    AACTGCGTGACCCATTCCGGATGGGGAACCTTAGGAATTAAAGCAATCTCGGTTCTATTTAACGTTTCCATAGACATGGTGTTGGAGAAAAAGTTGTTGACCAACCTGCAAACATCATTT
FR    ------------------------------------------------------------------------------------------------------------------------

SY    CCTACTATGCTCCAATACTTTTGGTAAAAAATTCCAGAGAAACCATCAGGTCCTGGCGATTTCAAGGCCCCCATTTGGAAAACAGTAATTCGAATTTCCTCATTTGAGATCGGGGCCAAA
FR    ------------------------------------------------------------------------------------------------------------------------

SY    AGAGAAGCATTAATGTCGTCAGAAATAACTACAGGAACGAAATCTAGAATATCCCCCCAATCTCTTGGCCCTTCGGACGTGAATAAGTTTTTAAAATAATCTTCGATTATACTCCGCACG
FR    ------------------------------------------------------------------------------------------------------------------------

SY    TGATGTTCACCCATTTCCCAGTCTCCAGCATGATTCTTTATTCTTTCCAGTCGATTCCTTCGCCTTTTTTGAATTGTGCAATTATGGAAAAAAGCAGTATTTGCATCTCCATGTTTTAGC
FR    ------------------------------------------------------------------------------------------------------------------------

SY    CACAGGGTTTTTGCCCTCTGCTTCCAGTACAACTCTTCGCACCTCCAAGTGTGGTTAAGGCTCCTCTCCACTTCCTTGATCTTCATGGTATTTTCCTCCCAGTCTTGTTGGAGGCTGTCT
FR    ------------------------------------------------------------------------------------------------------------------------

SY    AATTCAGATAGGAGAGCAGTGGCCATTTTCCGATTGTTTTGAAATTTTCCATCACTCCACTGTTTCAAATCATTTCTGCAGGTTCCTAACTTTGTGTCCCAGGAGAAGCGAGAGGCAGCA
FR    ------------------------------------------------------------------------------------------------------------------------

SY    GGAGTGCAAGGCCCCCAACTCCTATCAACTACCTCTCTGCATTCAGGATCAGATGCCCAGAACGCCTCAAATTTGAACGGCTTCAAACCCCTCCTAACATTTGTCTCAGTCAGAATCAGA
FR    ------------------------------------------------------------------------------------------------------------------------

SY    ACAGGACAATGATCCGAGCCCACAGCTGGTAAATGGATGGCATGGGAGTTTGGCCAAGATTCCTGCCAAGGCACATTTATCAGCCCACGGTCAAGTCTTTCCTGCACAACCACTCCGTCA
FR    ------------------------------------------------------------------------------------------------------------------------

SY    GCCCTAGTCCCTCTCCAAGTGAAAGACGAACCCTGATACCCCAGGTCCAACAGCTCTTTTTTATCTAAAAATTCTTGTAGATACCTTCTCCTGTTGTTGTCTAACCTCCTTCCACCTCTC
FR    ------------------------------------------------------------------------------------------------------------------------

SY    TTCTCAAAATCCCAAAGCATATCATTGAAATCACCAAAGCATAACCAAGGAAAATCCACAGACCCCAGGACTGAATCCAACCACCCCCAACAAGCCTCCTTCTCTTCCCGGTATGGAGAA
FR    ------------------------------------------------------------------------------------------------------------------------

SY    CCATAAACCCACGAAGCTCTAAATCTAACACCCGACCCTTGCTCAGTCACCCAAGAGTCAATCAGATTTTTTGAGCAATTAGTAACCTCCACTTCAACCCTGTCATCCCACCACAAGCAT
FR    ------------------------------------------------------------------------------------------------------------------------

SY    AAACCCCCAGATGTGCCAACTGGGTCCACGTAAAACTCATGAACAAACCCCACATCCCTTGCCAACCTTGTTAACCTATTTTGTTTCTGCTTCGTTTCCATGAGAAACACCATGGAGGGT
FR    ------------------------------------------------------------------------------------------------------------------------

SY    CGCTTTCGTTTTATTGTCTCCCGAAGAGACCGAATTGTCAGGTCCGACCCAATACCTTGACAATTCCATGCTAGGTAACTCATGGCGACCTTGTGGCTGTTTTTGGCCAGCCACCACCGC
FR    ------------------------------------------------------------------------------------------------------------------------
SY    CATTTTCTGCTGCACTTGTAACTTCAAATTGTGAGTCCCTCAACTCCAATGTCGGCGCCAAGACCACTGCCACATCCCTCAACCCTTCCTCTGAGAATTCAATTACCTGGGGGACCAGTC
FR    ------------------------------------------------------------------------------------------------------------------------

SY    CATGCGACTTCGTCGTCCTAGCTTTAGAACTTTTAGCGAACCTTCGTTGTTTATTCCTTCCAGTCTTCACCTCAGCAACTAGCCCTTCACCTTTCAAGACTTTTGGTTTTTTGCCTTCCT
FR    ------------------------------------------------------------------------------------------------------------------------

SY    TCAAAGATCTCCAGCTGGACTCAGCTTCTCTCTTAACGCCCATTCTTTCAAAAGCAGCATTGGCATTTAAACTAGGCCCATGAATTGAATCAGTATGGTTTTGTCCAACATGATTTGTAG
FR    ------------------------------------------------------------------------------------------------------------------------

SY    GGCTTAGATCCTTCCCAGGTTGGGAATCCAACTCCTCAAGAGCTACAGAATTTCGGCCCTTCTGTGAAGGCCCAAGCAAGGCCCAATTTGGAGCAGCATGCTGACCAAAAGACATTTGAG
FR    ------------------------------------------------------------------------------------------------------------------------

SY    AGTGGGCCCATTGAATTTTATCCCAGTGGCGTGAATTATGCTCCTCAACTAGAATCACCCCACTCCGACCTAGAGAGCTAGTACTTCCCCTGACACTATTAGTTTGCCACTCAAAAATCT
FR    ------------------------------------------------------------------------------------------------------------------------

SY    GCATGTGAGGGTGTGAAACCTGTCTACCGCCCACTTCTCTTGCGCTATCACTTTCAATCCTTTCCTCCTGACCAGGGGATTGATTTGAAATAGCCGTTTGCCCCATTATTTTATTGCATA
FR    ------------------------------------------------------------------------------------------------------------------------

SY    TAGGCTCAGTCAGTTGTTGTCTCGAGGTTTCTTGGTTTGTGTGTGCAACGTCTCCATCAGCTTCCTCCCTATCTCCACCTTCCGTGTGCAAGTGGCCCACACGCGCCACCCTAGGAACCA
FR    ------------------------------------------------------------------------------------------------------------------------

SY    GGTTGGCGATACTTCTTTCCCCTGCTTTACGACGGACAGCCCGAGGGGTAATGGAGTTACTCTGGCCTCGCCTTGCCCGAATAGCTCTCGCTTTCATCCAGTCACCATAAGCAGCCGCTT
FR    ------------------------------------------------------------------------------------------------------------------------

SY    CCGCACCCTCCTCCTGATGCGCCAGAACTTTGCAGCCGTTCTTCGTGTGACCCAGCCTCCCACAGTCATAGCAAAAATCGGATAGCCTCTCATACTTCAATTCCACCCATGTGGCCAGGC
FR    ------------------------------------------------------------------------------------------------------------------------

SY    CTTCGGCTCTTGGGAGCCAAAACCCAGCAGCTAAGGGGGCTCGACAATCGATCTGGATTCGGATCCTCAAGAAGCCTCGCGGACCCATGCTCCAAGGATCTTCAACTTCCAGAACAGATC
FR    ------------------------------------------------------------------------------------------------------------------------

SY    CAAGCTTTCCACCAATCTCCGTTGCGTTCTCGGTCGACATGAGGTTAAGAGGTATTCCATGAATTTGGATCCAATAAGCAACAAGGTGAGTCGGAATTTCTTCAATCGCCATGCTACTTG
FR    ------------------------------------------------------------------------------------------------------------------------

SY    GCCATCTCTGCACTGAAAAGCAAAAGCCCATTACTGACCAAGGGCCCTCGTCAACAATTTTATCCGCCGTATTTTCATCCTGGGCAAGGATAGTAAACACATTCTCCTTCATGTATGTGA
FR    ------------------------------------------------------------------------------------------------------------------------

SY    TTTTCGTCTCCCCAAAGGAAGCCCAGGCTGTCGACAGGATGTTCTTGATAGCACCCCTATTTGGAATCTTGTCAGCAACGATGGCACCAATCAATTTTGTTCCACTCTCCAGGCTTGAGA
FR    ------------------------------------------------------------------------------------------------------------------------

SY    AGCCAAGGGCCTCTTCAAGTTTAATAACCAGCTCCTCCGTTCTAGCGTCCGCCATAGTGGTTTCCAAAGATGACCCGCACGTGTAGTTTTCTATCAAGTAAACAGGTTCTGTTTGGGTTG
FR    ------------------------------------------------------------------------------------------------------------------------

SY    GGTTTGTAGGAAACACTATCAGAGTTTGTTTCCAGTGAAAATTCTTTTGCAGGGGGGTAGTTTCAGTTCTATTTGGGCTCCAGGGAAAGATTGTATGGTTGTTGTCTTGATGGTAGTAGG
FR    ------------------------------------------------------------------------------------------------------------------------

SY    TGGCCCAAGATCTGTACCAAACTAGGGTCTTTATTACCCAGCTCATCCAGTCTTGTTTTGTGCTCATAACTCTCCTCTTCTTCGATATCTGGGAGGAGTAAATGTCTGCCGCCCTTCTAC
FR    ------------------------------------------------------------------------------------------------------------------------

SY    CAGAGAGGTAGGTGAGAGTCATAACTCCAAGCTCCCAAGGAAGCTTGTTACTGTGAACATTTCGTTGAGGGAAAAATGACTCACACATAGGTGAGGGAACTCTCATTGTAGAGAGAGAGG
FR    ------------------------------------------------------------------------------------------------------------------------

SY    CTCTCATTGTAGAGAGAGGCTGAACTATATGCCAACATGATTATGAGTGAGATTATGGCACTGGAGAAAAAGGTTAGATGATTGATACGTACTCTGTAAATGAAAACAAATTTCGACTAT
FR    ------------------------------------------ATGAGTGAGATTATGGCACTGGAGAAAAAGGTTAGATGATTGATACGTACTCTGTAAATGAAAACAAATTTCAACTAT

SY    TTACAATAATCAGGACTTTAACATCAATAATCATAACCAAGTTTGGCTCTGGTTTTCTCTTCCACGCACACCGACTATAATTTCAGGGAGCTGAGCTGGTAGAAGCCAACAACCAGTTGA
FR    TTACAATAATCAGGACTTTAACATCAATAATCATAACCAAGTTTGGCTCTGGTTTTCTCTTCCACGCACACCGACTATAATTTCAGGGAGCTGAGCTGGTAGAAGCCAACAACCAGTTGA

SY    GGCAGAGGGTAAGCAACTACCACAATCATGTACATATTTCTTCTTCTTTTTTTATTTTTATTTTGTATTTTTTTGCATAGCAGATGGTGATGTTATCCGGAGGAAATACTGGACCTGCGT
FR    GGCAGAGGGTAAGCAACTACCACAATCATGTACATATTTCTTCTTCTTTTTTTATTTTTATTTTGTATTTTTTTGCATAGCAGATGGTGATGTTATCCGGAGGAAATACTGGACCTGCGA

SY    TTGTGGAGCCGGAGACGTTGATTACTAATGTTGGAGGTGGAGGAGAAGAAGACGGCATGTCATCTGAATCTGCCCTAATTGCCACCTCCACCAGCTGCAACAGTGATGTCAGTCTCTCTC
FR    GTGTGGAGCCGGAGACTTTGATTACTAATGTTGGAGGTGGAGGAGAAGAAGACGGCATGTCATCTGAATCTGCCCTAATTGCCACCTCCACCAGCTGCAACAGTGATGTCAGTCTCTCTC

SY    TTGAAGATGACTGCTCCAATGTCACTTTATCTCTCAAATTGGGGTGAGCTACTTTTGCTTTGCTACGTATATTTATATTATATATTTATATGTTTTTTTTTTTATAGTTTGATCAGAGAA
FR    TTGAAGATGACTGCTCCAATGTCACTTTATCTCTCAAATTGGGGTGAGCTACTTTTGCTTTGCTACGTATATTTATATTATATATTTATATGTTTTTTTTTT-ATAGTTTGATCAGAGAA

SY    AACATGCTTGCTTTATTTGATGATATTATAAAGAATAAATTAAAAGTT------------------------------------------------------------------------
FR    AACATGCTTGCTTTATTTGATGATATTATAAAGAATAAATTAAAAGTTAAAAAAACACAACTGTATGGGAGATCTAGCGCCGCTCTCCCAAACAACCAAACCCTAGATCTAGCGCCGCTC

SY    ------------------------------------------------------------------------------------------------------------------------
FR    CATATCTGCGAGGGGGAGGAGGAGTCCTCCTCCCCCCTCCCTCTCTACTGGATCTTGCTAGCCTTTCGACGATTCTGTCTCGGAGCATTTGGAATCCCCATCCGTAGCTCTCTCTTTTGA

SY    ------------------------------------------------------------------------------------------------------------------------
FR    GGGCTGGTGACGTGTTTCGAGCTTCACCTGCTCGTTTTTGTGGCATCATGCCCCCTTCCTTGTTTGCATCCCCAAATCTTCATTATACCTTTCCCCTCAGAGATGTCTTTTGGCTATCCC

SY    ------------------------------------------------------------------------------------------------------------------------
FR    CCGATTGGCTGTTTTTATCTTCTAGGTGCGCCGATTTCGAATCTGGTGGAGCGGGGGTCCCTCACTCTCCCATCTCGACCTGGTTGTGTCTATTGGATATTGCAGCTAATTCGAAACGCG

SY    ------------------------------------------------------------------------------------------------------------------------
FR    ATCTCTCGCGTAAATCTTGGAAGATGTGTAGAGCCACGGCAGTGCTACGGCTCGGGTGCTTGCACCACCTCCTTCCTTTTTCATGGTTCCGGTTCTCTCGGCTTTTGACGGATTTGAGGG

SY    ------------------------------------------------------------------------------------------------------------------------
FR    TTTTGGTTATGGTGGTGGATCTGTGGCTGATAAGCTTATCATTTGTTTGTTGTTCAATTGGGCATTTTGTTGTTTTATTTCAAATTGTTGTTGTTTGTTGTAACTTCTGTTTATCTGTTG

SY    ------------------------------------------------------------------------------------------------------------------------
FR    GTATCAACATCTGGGACAATCTTGGTGTTCTTGTTCATGTGATATCACAGATTAGTTATTCTGTTGTGCTTCCTTACTGGATTGCTGATGATGTAATTCTGGGCTCTCCCCGTTTTGGGT

SY    --------------------------------------------------------------------GGATCTCGTGCAAGAAGAGGGTTTTGCTTAGATATATTTCTGTACTTGTCAA
FR    GAGGTTCTTGTTTTATGAAATATCAACTTTCGACCAAAAAAAAAAAAAAGTTAAAATAATCAATACTTGGATCTCGTGCAAGAAGAGGGTTTTGCTTAGATATATTTCTGTACTTGTCAA

SY    CTTGATTTAGGGTTTCATTTCATATGAACTCTACCAAACCAAAATATTTGGATTATTAGATTGAAGATGTTCTAGGAGTTTGTAGAGCAAGTTTATTTTGAGGTTTCATGTTTGTGTCTT
FR    CTTGATTTAGGGTTTCATTTCATATGAACTCTACCAAACCAAAATATTTGGATTATTAGATTGAAGATGTTCTACGAGTTTGTAGAGCAAGTTTATTTTGAGGTTTCATGTTTGTGTCTT

SY    TGGGCTATTTCATTTTTGTGTTTATTTAAATTGAAACAAGTTAGTGACTTTCATCGAAACTCTCTCACATTTTTTTTATATTTTTTGTTATAGAAAATATTTATATAAAAAATGTGAATT
FR    TGGGCTATTTCATTTTTGTGTTTATTTAAATTGAAACAAGTTAGTGACTTTCATCGAAACTCTCTCACATTTTTTTTATATTTTTTGTTATAGAAAATATTTATATAAAAAATGTGAATT

SY    TCGTAGTATGGCAGTCCAAATAAAATTATTTTAAATATTTTTATATAACTAATAATAATTAACTTGCTACAGAAATCTTCAGTTACGAAATCCCAACACTAA------------------
FR    TCGTAGTATGGCAGTCCAAATAAAATTATTTTAAATATTTTTATATAACTAATAATAATTAACTTGCTACAGAAATCTTCAGTTACGAAATCCCAACACTAATGGAGAAACTAAGGAATC

SY    ------------------------------------------------------------------------------------------------------------------------
FR    AGAGAATTTAGGAATCGGAGAACTACTGAATCGGTATGGGAAGAATCATTCCCATTAAGCTGTTTACTAAACATATGGAATTAGCAATGGAATGTGATTGATTCCCATTGTTATTGTTTA

SY    ------------------------------------------------------------------------------------------------------------------------
FR    CTAATTTGCATGAATCAAAATCCAAATTAATTTTAGAATTATGGTAAAGTGATAAGAGTATTTTTGGAAGTTATTAAGAGAAAACAGGAATGTGAATCGTATCGACTACTCCCCCTTAGT

SY    ------------------------------------------------------------------------------------------------------------------------
FR    TGATCTGATTCCTAGTTTTGAGGGAATGTGATTACGGATTTTGAGGTGGGGCTCACTTATTTTTTCATTCCTGATTGCAAGTAAACGCAGGAGAAGTCAGGAATGGAAATCATTCCCTTT

SY    --------------------------------------AAGGGCTATTGGAAACATACCGGAGAAAATATATTCCACTCTCTCTAGACTAAAACAATGTAACCTGAATAAGGCATATAAA
FR    CCTCTCATTCCCATTACTGATACGTAAACATGCCCTAAAAGGGCTATTGGAAACATACCGGAGAAAATATATTCCACTCTCTCTAGACTAAAACAATGTATCCTGAATAAGGCATATAAA

    SY    GGTTAAAATATCTCCAACAAATGTTGATGGGCAATTGCATTCATGGTATTTCATTTAATTCTTCACCTTTTTAATCAATTTGCTAATCACTTTGTATTTTCTCTGAATTTCAGGCTTCCCTAG
    FR    GGTTAAATTATCTCCAACAAATGTTGATGGGCAATTGCATTCATGGTATTTCATTTAATTCTTCACCTTTTTAATCAATTTGCTAATCACTTTGTATTTTCTCTGAATTTCAGGCTTCCCTAG
```
